# A comprehensive score reflecting memory-related fMRI activations and deactivations as potential biomarker for neurocognitive aging

**DOI:** 10.1101/2021.01.16.426666

**Authors:** Joram Soch, Anni Richter, Hartmut Schütze, Jasmin M. Kizilirmak, Anne Assmann, Gusalija Behnisch, Hannah Feldhoff, Larissa Fischer, Julius Heil, Lea Knopf, Christian Merkel, Matthias Raschick, Clara-Johanna Schietke, Annika Schult, Constanze I. Seidenbecher, Renat Yakupov, Gabriel Ziegler, Jens Wiltfang, Emrah Düzel, Björn H. Schott

## Abstract

Older adults and particularly those at risk for developing dementia typically show a decline in episodic memory performance, which has been associated with altered memory network activity detectable via functional magnetic resonance imaging (fMRI). To quantify the degree of these alterations, a score has been developed as a putative imaging biomarker for successful aging in memory for older adults (*Functional Activity Deviations during Encoding*, FADE; Düzel et al., 2011). Here, we introduce and validate a more comprehensive version of the FADE score, termed FADE-SAME (*Similarity of Activations during Memory Encoding*), which differs from the original FADE score by considering not only activations but also deactivations in fMRI contrasts of stimulus novelty and successful encoding, and by taking into account the variance of young adults’ activations. We computed both scores for novelty and subsequent memory contrasts in a cohort of 217 healthy adults, including 106 young and 111 older participants, as well as a replication cohort of 117 young subjects. We further tested the stability and generalizability of both scores by controlling for different MR scanners and gender, as well as by using different data sets of young adults as reference samples. Both scores showed robust age-group-related differences for the subsequent memory contrast, and the FADE-SAME score additionally exhibited age-group-related differences for the novelty contrast. Furthermore, both scores correlate with behavioral measures of cognitive aging, namely memory performance. Taken together, our results suggest that single-value scores of memory-related fMRI responses may constitute promising biomarkers for quantifying neurocognitive aging.

## 1. Introduction

In functional magnetic resonance imaging (fMRI) studies of episodic memory, a widely used approach is to probe the successful acquisition of novel information (*encoding*) as a function of performance in a later memory test (*retrieval*), the so-called “subsequent memory effect” or difference due to later memory (DM) effect (Paller et al., 1987). Since its first application to fMRI (Brewer, 1998; Wagner et al., 1998), numerous studies have employed this approach, and meta-analytic evidence shows that successful encoding robustly engages the medial temporal lobe (MTL) as well as inferior temporal, prefrontal, and parietal cortices (Kim, 2011). When compared to young adults, older individuals display characteristic differences in memory-related network activations, including a reduced activation in the MTL, particularly the parahippocampal cortex and a reduced deactivation or even atypical activation of midline cortical structures (Düzel et al., 2011; for a review and meta-analysis see Maillet and Rajah, 2014). While such functional age-related differences are often accompanied by cognitive decline, as evidenced, for example, by lower later memory performance or poorer recollection of episodic details (Cansino, 2009; Wong et al., 2012), age-related differences in memory-related brain activity do not necessarily indicate worse memory performance. Instead, differences in functional neural networks may also reflect adaptive strategies that are employed as a compensatory mechanism and can even be accompanied by better behavioral performance (Cabeza et al., 2002, 2018; Grady & Craik, 2000; Stern, 2009). Early fMRI studies have shown that older adults who did not differ in their episodic memory performance from young adults exhibited increased recruitment of prefrontal cortex (PFC)-dependent neurocognitive resources (Cabeza et al., 2002; Grady & Craik, 2000). Such findings highly suggest that at least some older adults can draw from some sort of cognitive and/or neural reserve. Theories of neural reserve (i.e., individual quantitative or qualitative differences of brain anatomy that underlie the efficacy of neural networks) and cognitive reserve (individual approaches of processing tasks) propose that the time point at which behavioral performance shows decline heavily depends on the outset level of an individual and, thus, their ability to compensate age-related physiological alterations (e.g., Stern, 2009).

Despite this well-documented inter-individual variability of cognitive and neural aging, age-related differences in the neuroanatomical underpinnings of successful episodic encoding are rather robust at the group level (Cansino, 2009; Maillet & Rajah, 2014). Considering the well-replicated observation of – on average – lower memory performance in older adults, it seems worthwhile to explore the value of such functional deviations of episodic memory encoding networks from young participants – concurrent with the assessment of behavioral memory performance – as a potential tool for the quantification of neurocognitive aging. However, few studies have explicitly tested the applicability of age-related differences in memory-related fMRI activations as an individual biomarker for cognitive aging. One such approach assessed the influence of a genetic risk factor for accelerated neurocognitive aging and Alzheimer’s disease, apolipoprotein E (ApoE), on structural and functional MRI indices of age-related memory impairment (Woodard et al., 2010). The authors compared the predictive value of different combinations of ApoE genotype (ε4 allele present/not present), hippocampal volume, and fMRI novelty response (recognition of famous vs. unfamiliar faces), and concluded that a combination of ApoE genotype and fMRI novelty response were the best predictor of cognitive decline, enabling the correct classification of 78.9 % of all participants. While such approaches provide important insight into the factors contributing to cognitive aging, their clinical application is often limited. Typically, clinicians want to use single scores derived for a certain individual to decide, based on comparison to normative thresholds, how to classify that person and, for example, their likelihood to develop Alzheimer’s disease in the near future.

There are only very few studies that evaluated the use of single-value scores for assessing age-related differences of functional memory networks.^1^ One such study (Salami et al., 2012) used a multivariate statistical approach, partial-least-squares (PLS), to extract latent variables that covaried with fMRI activations related to a face-name-association encoding task, a corresponding retrieval task, and a baseline perceptual change-detection task. The authors found that the degree of engagement of the encoding/retrieval network was predictive of age-related memory performance. However, it cannot be excluded that such a latent variable may be highly task-dependent and not sufficiently generalizable to prototypical functional episodic memory network activity.

It is thus desirable to develop a reductionist score that can be applied within the known reference framework of age-related differences in memory-related network activity. To the best of our knowledge, the only study that actually tested the utility of a reductionist fMRI-based biomarker within such a reference framework was a study conducted by Düzel et al. (2011). They proposed a single-value score in which the age-related differences of encoding-related network activations are described in a single number that denotes the degree of deviation from the prototypical activation pattern observed in young adults (*Functional Activity Deviation during Encoding*, FADE; Düzel et al., 2011). In the original study by Düzel and colleagues (2011), the FADE score was based on neural correlates of successful memory encoding, namely, the DM effect, but this approach may be limited in participants with poor memory performance, due to lack of successfully encoded items (Soch et al., 2021). To circumvent this limitation, one might base the FADE score calculation on the novelty effect, namely the brain’s response to novel information, irrespective of encoding success, an approach supported by recent observations that hippocampal novelty responses correlate with tau protein concentrations in cerebrospinal fluid (CSF) in older adults (Düzel et al., 2018). An alternative, or perhaps complementary, approach to focusing may be the use of a parametric model of the DM effect, which can also be computed in individuals with relatively poor memory performance (Soch et al., 2021). However, both approaches have not yet been used in the context of the FADE score and therefore warrant validation.

The aim of the present study was two-fold: On the one hand, we aimed to validate the use of a single numeric value reflecting memory-related fMRI activation differences as a proxy of cognitive aging in a large cohort of healthy older participants, using both the novelty contrast and a parametric DM effect. Secondly, we aimed to extend the original FADE score (hence termed FADE-classic) by (i) considering both activations and deactivations during encoding and by (ii) taking into account the variance of the reference sample of young subjects required for its computation, thereby yielding the so-called FADE-SAME score (*Similarity of Activations during Memory Encoding*). Specifically, the following features were implemented:

i. Both novelty and DM contrasts engage a similar set of brain regions, including the MTL with the parahippocampal cortex and hippocampus, inferior temporo-occipital and lateral parietal cortices, and the dorsolateral prefrontal cortex (dlPFC) (Düzel et al., 2011; Soch et al., 2021). Collectively, these brain regions can be considered to constitute a human memory network. Notably, older subjects do not only show reduced activations in this network, particularly in the parahippocampal cortex, but also reduced *de*activations in brain regions like the ventral precuneus and posterior cingulate cortex (PreCun/PCC; see Figure 2), which are part of the brain’s default mode network (DMN) (Maillet & Rajah, 2014; S. L. Miller et al., 2008). These deactivations were already mentioned in the original study of the FADE score (Düzel et al., 2011) and are now explicitly considered when computing the FADE-SAME score.
ii. The FADE score reflects, by definition, the deviation of an older adult’s memory-related activation pattern from the prototypical pattern seen in young adults. It therefore requires referencing the activation map of the respective individual to a baseline activation map obtained from a cohort of young adults (Düzel et al., 2011). However, memory-related fMRI activation patterns also exhibit individual differences among young adults, which are stable over time and thus likely reflect traits (M. B. Miller et al., 2002). To avoid potential biases related to individual activation patterns of the specific young adults contributing to the FADE score template, it is thus advisable to account for the variance of the reference sample itself, which is implemented in the calculation of the FADE-SAME score.

To evaluate how both the classic FADE score and the FADE-SAME score perform as potential biomarkers of neurocognitive aging, we compared the two scores in a large sample of healthy young (N = 106; age range: 18-35 years) and older participants (N = 111; age range: 60-80 years) studied within the *Autonomy in Old Age* project^2^ (Assmann et al., 2020; Soch et al., 2021). This study used a shortened version of the subsequent memory paradigm from the original FADE study (Düzel et al., 2011), which is also employed in a large-scale longitudinal study of pre-clinical stages of Alzheimer’s disease (Bainbridge et al., 2019; Düzel et al., 2018). In this paradigm, photographs of scenes are encoded incidentally via an indoor/outdoor decision task, and memory is tested via an old/new recognition memory task with a five-step confidence rating. The FADE-classic and FADE-SAME scores were computed on activation maps from both successful memory encoding (Düzel et al., 2011; Soch et al., 2021) and novelty processing (Düzel et al., 2018), and evaluated with respect to their power to differentiate between age groups and their correlation with memory performance and hippocampal volumes.

## 2. Methods

### 2.1. Participants

The study was approved by the Ethics Committee of the Otto von Guericke University Magdeburg, Faculty of Medicine, and written informed consent was obtained from all participants in accordance with the Declaration of Helsinki (World Medical Association, 2013).

Participants were recruited via flyers at the local universities (mainly the young subjects), advertisements in local newspapers (mainly the older participants) and during public outreach events of the institute (e.g., *Long Night of the Sciences*).

The study cohort consisted of a total of 217 neurologically and psychiatrically healthy adults (see Table 1), including 106 young (47 male, 59 female, age range 18-35, mean age 24.12 ± 4.00 years) and 111 older (46 male, 65 female, age range 60-80, mean age 67.28 ± 4.65 years) participants. According to self-report, all participants were right-handed, had fluent German language skills and did not use neurological or psychiatric medication. The Mini-International Neuropsychiatric Interview (M.I.N.I.; Sheehan et al., 1998; German version by Ackenheil et al., 1999) was used to exclude present or past psychiatric illness, alcohol or drug dependence. Age groups were not significantly different with respect to ApoE genotype (see Table 1; also see Supplementary Figure S3A), but there were differences regarding medication (see Supplementary Table S2) and with respect to educational years: While 94% of young subjects received the German equivalent of a high school graduation certificate (“Abitur”), this was only the case for 50% of the older subjects, most likely due to historical differences in educational systems (see Supplementary Discussion for potential explanations of the demographic between-group differences). Using a multiple-choice vocabulary-based screening of verbal intelligence (“Mehrfachwahl-Wortschatz-Intelligenztest”, MWT-B; Lehrl, 2005), we could confirm that older participants had comparable or even superior verbal knowledge (see Table 1).

**Table 1.** Demographics of young and older subjects.

|  | **young subjects** | **older subjects** | **statistics** |
| --- | --- | --- | --- |
| N | 106 | 111 | – |
| age range | 18-35 yrs | 60-80 yrs | – |
| mean age ± SD | 24.12 ± 4.00 yrs | 67.28 ± 4.65 yrs | $t = -73.11, p < 0.001$ |
| gender ratio | 47/59 m/f | 46/65 m/f | $\chi^2 = 0.19, p = 0.666$ |
| ethnic composition | 104/2<br>European / other | 111/0<br>European / other | $\chi^2 = 2.11, p = 0.146$ |
| educational status | 100/6<br>with / without Abitur | 56/55<br>with / without Abitur | $\chi^2 = 51.68, p < 0.001$ |
| ApoE genotype | 1/15/4/60/24/2<br>E2/E2 / E2/E3 / E2/E4 /<br>E3/E3 / E3/E4 / E4/E4 | 2/20/4/69/16/0<br>E2/E2 / E2/E3 / E2/E4 /<br>E3/E3 / E3/E4 / E4/E4 | $\chi^2 = 5.16, p = 0.396$ |
| mean MMSE<br>performance ± SD | – | 28.84 ± 1.02<br>(range: 26-30) | – |
| MWT-B hits ± SD | 26.81 ± 3.14 | 30.60 ± 3.00 | $z = -8.15, p < 0.001$ |

Demographic information for the two age groups, along with statistics from a two-sample t-test (mean age), chi-squared tests (gender ratio, ethnic composition and ApoE genotype) and a Mann-Whitney U-test (MWTB hits). Abbreviations: N = sample size, SD = standard deviation, yrs = years, m = male, f = female, MMSE = Mini-Mental State Examination (Creavin et al., 2016; Folstein et al., 1975), MWT-B = multiple choice vocabulary intelligence test (“Mehrfachwahl-Wortschatz-Intelligenztest”; Lehrl, 2005). “Abitur” is the German equivalent of a high school graduation certificate qualifying for academic education.

### 2.2. Experimental paradigm

During the fMRI experiment, participants performed a visual memory encoding paradigm with an indoor/outdoor judgment as the incidental encoding task. Compared to earlier publications of this paradigm (Assmann et al., 2020; Barman et al., 2014; Düzel et al., 2011; Schott et al., 2014), the trial timings had been adapted as part of the DZNE-Longitudinal Cognitive Impairment and Dementia (DELCODE) study protocol (see Bainbridge et al., 2019; Düzel et al., 2018; Soch et al., 2021, for a detailed comparison of trial timings and acquisition parameters). Subjects viewed photographs showing indoor and outdoor scenes, which were either novel at the time of presentation (44 indoor and 44 outdoor scenes) or were repetitions of two highly familiar “master” images (22 indoor and 22 outdoor trials), one indoor and one outdoor scene pre-familiarized before the actual experiment (cf. Soch et al., 2021, Fig. 1B). Thus, every subject was presented with 88 unique images and 2 master images that were presented 22 times each. Participants were instructed to categorize images as “indoor” or “outdoor” via button press. Each picture was presented for 2.5 s, followed by a variable delay between 0.70 s and 2.65 s. To optimize estimation of the condition-specific BOLD responses despite the short delay, simulations were employed to optimize the trial order and jitter, as described previously (Düzel et al., 2011; Hinrichs et al., 2000).

**Figure 1.**
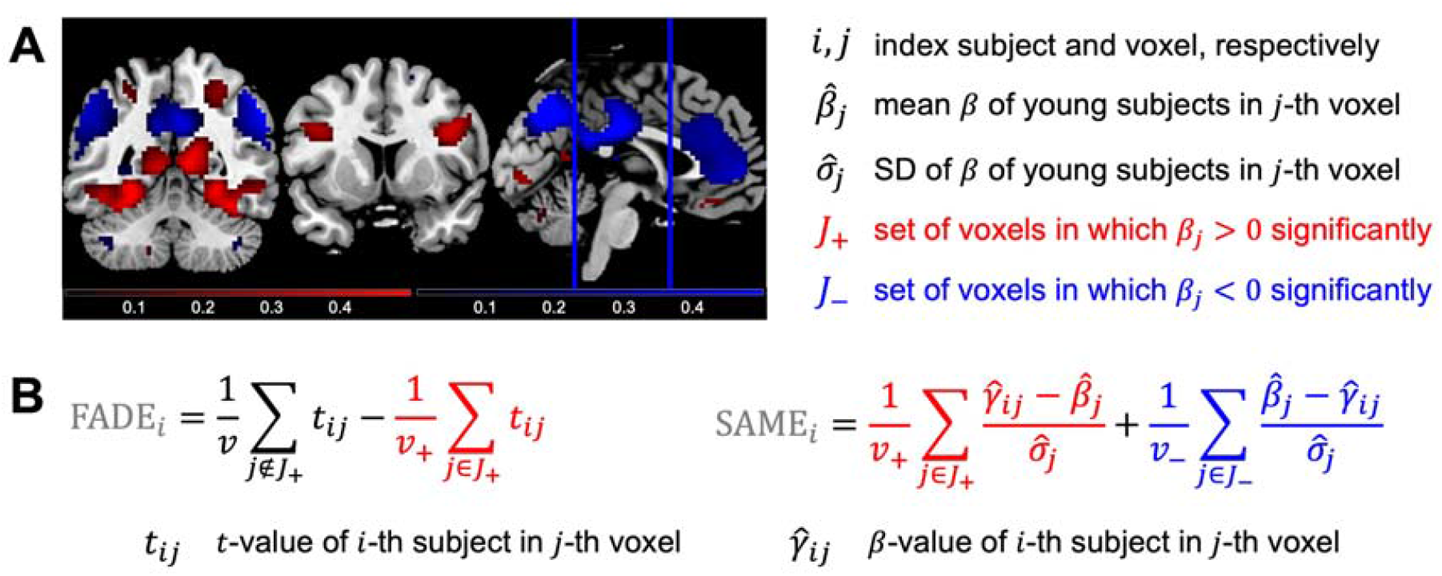
Measures for quantifying successful aging in memory. We compute two summary statistics from fMRI contrasts, which are both based on a group-level analysis across all young subjects and subject-wise computation in each older subject. **(A)** A reference map is obtained by significance testing of a contrast within the group of young subjects, resulting in voxels with significant activation (red) or significant deactivation (blue). **(B)** FADE-classic and FADE-SAME score of older subjects are calculated as summary statistics by averaging single-subject contrast outcomes within selected sets of voxels (for explanations, see text).

Approximately 70 minutes (70.19 ± 3.60 min) after the start of the fMRI session, subjects performed a computer-based recognition memory test outside the scanner, in which they were presented with the 88 images that were shown once during the fMRI encoding phase (*old*) and 44 images they had not seen before (*new*). Participants rated each image on a five-point Likert scale from 1 (“definitely new”) to 5 (“definitely old”). For detailed experimental procedure, see Assmann et al. (2020) and Soch et al.(2021).

### 2.3. fMRI data acquisition

Structural and functional MRI data were acquired on two Siemens 3T MR tomographs (Siemens Verio: 58 young, 64 older; Siemens Skyra: 48 young, 47 older), following the exact same protocol used in the DELCODE study (Düzel et al., 2019; Jessen et al., 2018).

A T1-weighted MPRAGE image (TR = 2.5 s, TE = 4.37 ms, flip-α = 7°; 192 slices, 256 × 256 in-plane resolution, voxel size = 1 × 1 × 1 mm) was acquired for co-registration and improved spatial normalization. Phase and magnitude fieldmap images were acquired to improve correction for artifacts resulting from magnetic field inhomogeneities (*unwarping*, see below). For functional MRI (fMRI), 206 T2*-weighted echo-planar images (TR = 2.58 s, TE = 30 ms, flip-α = 80°; 47 slices, 64 × 64 in-plane resolution, voxel size = 3.5 × 3.5 × 3.5 mm) were acquired in interleaved-ascending slice order (1, 3, …, 47, 2, 4, …, 46). The total scanning time during the task-based fMRI session was approximately 530 s. The complete study protocol also included a T2-weighted MR image in perpendicular orientation to the hippocampal axis (TR = 3.5 s, TE = 350 ms, 64 slices, voxel size = 0.5 × 0.5 × 1.5 mm) for optimized segmentation of the hippocampus (see Appendix C) as well as resting-state fMRI (rs-fMRI) and additional structural imaging not used in the analyses reported here.

### 2.4. fMRI data preprocessing

Data preprocessing was performed using Statistical Parametric Mapping (SPM12; Wellcome Trust Center for Neuroimaging, University College London, London, UK). EPIs were corrected for acquisition time delay (*slice timing*), head motion (*realignment*) and magnetic field inhomogeneities (*unwarping*), using voxel-displacement maps (VDMs) derived from the fieldmaps. The MPRAGE image was spatially co-registered to the mean unwarped image and *segmented* into six tissue types, using the unified segmentation and normalization algorithm implemented in SPM12. The resulting forward deformation parameters were used to *normalize unwarped* EPIs into a standard stereotactic reference frame (Montreal Neurological Institute, MNI; voxel size = 3 × 3 × 3 mm). Normalized images were spatially *smoothed* using an isotropic Gaussian kernel of 6 mm full width at half maximum (FWHM).

### 2.5. General linear modelling

For first-level fMRI data analysis, which was also performed in SPM12, we used a parametric general linear model (GLM) of the subsequent memory effect that has recently been demonstrated to outperform the thus far more commonly employed categorical models of the fMRI subsequent memory effect (Soch et al., 2021).

This model included two onset regressors, one for novel images at the time of presentation (“novelty regressor”) and one for presentations of the two pre-familiarized images (“master regressor”). Both regressors were created as short box-car stimulus functions with an event duration of 2.5 s, convolved with the canonical hemodynamic response function, as implemented in SPM12.

The regressor reflecting subsequent memory performance was obtained by parametrically modulating the novelty regressor with a function describing subsequent memory report. Specifically, the parametric modulator (PM) was given by

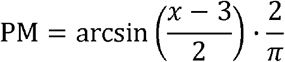

where *x* ∈ {1,2,3,4,5} is the subsequent memory report, such that −1 ≤ PM ≤ + 1. Compared to a linear-parametric model, this transformation puts a higher weight on definitely remembered (5) or forgotten (1) items compared with probably remembered (4) or forgotten (2) items (Soch et al., 2021, Fig. 2A).

**Figure 2.**
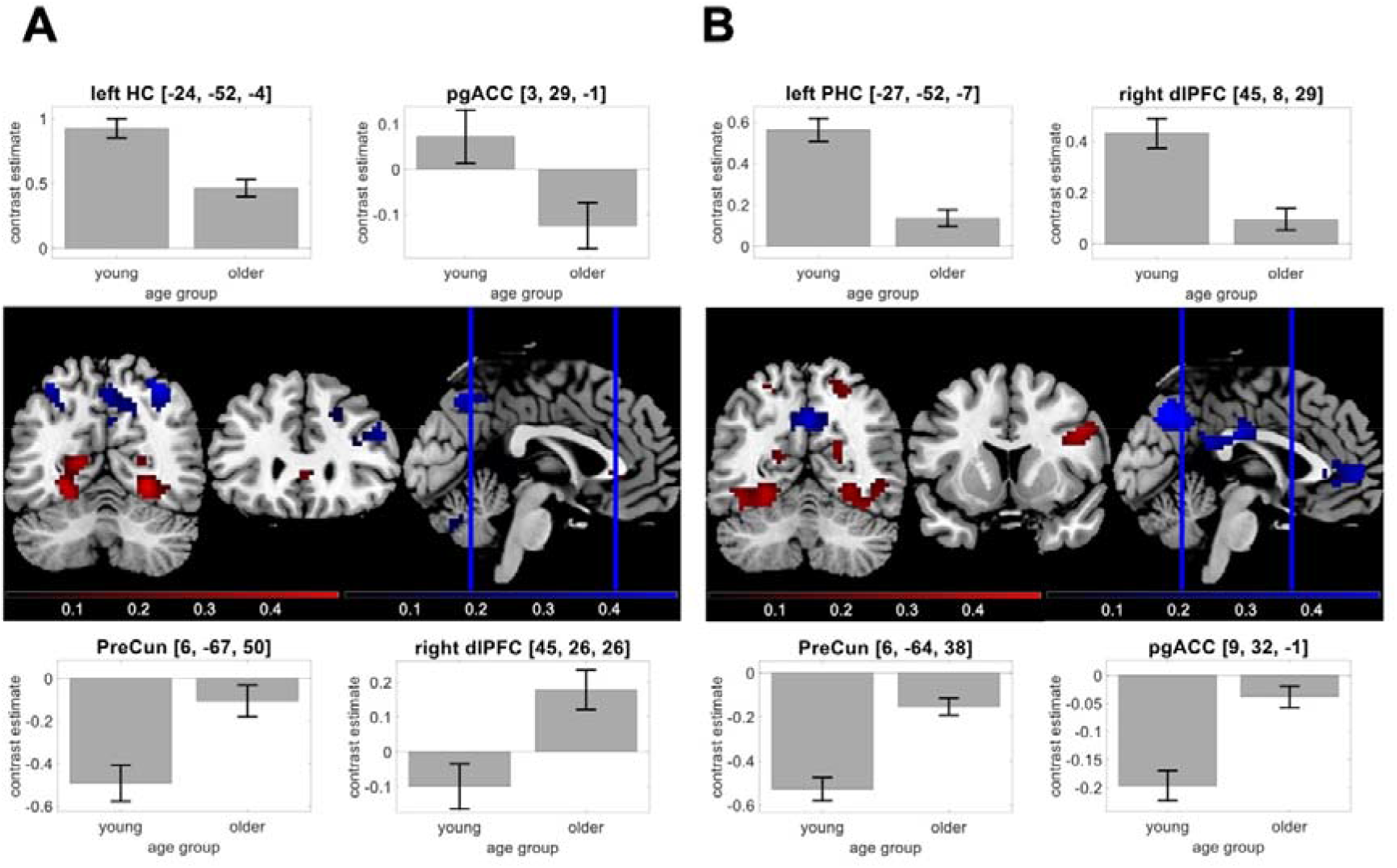
Age-related differences in the human memory network. Using our fMRI memory paradigm, we assessed novelty contrast and memory contrast and compared them between young and older adults. Brain sections show significant differences for activations (red) and deactivations (blue) in young subjects. Bar plots show group-level contrast estimates (gray) and 90% confidence intervals. **(A)** Significant effects of age on the novelty contrast, with reduced activations in hippocampus and pgACC and reduced deactivations in PreCun and dlPFC. **(B)** Significant effects of age on the memory contrast, with reduced activations in parahippocampal cortex and dlPFC and reduced deactivations in PreCun and pgACC.

The model also included the six rigid-body movement parameters obtained from realignment as covariates of no interest and a constant representing the implicit baseline.

### 2.6. Functional activity deviation during encoding (FADE-classic)

The original FADE score^3^ (here: FADE-classic) constitutes the first implementation of a single-value score of encoding-related fMRI activations designed as a potential biomarker in age-related memory decline. Computation of classic FADE scores canonically proceeds in two steps (Düzel et al., 2011, p. 805) (see Figure 2):

1. First, a reference map is generated by submitting contrast maps from young subjects to a group-level analysis and determining the set of voxels in which there is a significant positive effect (e.g., memory contrast: higher activations for items later remembered vs. later forgotten), with the entire set of voxels considered a “volume of interest” (VOI).
2. Then, the same contrast is computed for each older subject, resulting in a t-value map for each subject. Finally, the FADE score is obtained by subtracting the average t-value inside the VOI from the average t-value outside the VOI.

More precisely, let *J*_+_ be the set of voxels showing a positive effect in young subjects at an *a priori* defined significance level (p < 0.05, FWE-corrected, extent threshold k = 10 voxels in the present work^4^), and let *t* _*ij*_ be the t-value of the *i*-th older subject in the *j*-th voxel. Then, the FADE score of this subject is given by

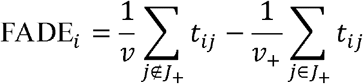

where *ν* _+_ and *ν* is the number of voxels inside and outside *J*_+_, respectively (see Figure 1B). While originally developed for the subsequent memory contrast (termed “recognition-encoding contrast” in the original publication), it is in principle also possible to calculate the score for the novelty contrast (novel images vs. familiar images). In either case, a larger FADE score signifies higher deviation of an older adult’s memory – or novelty – response from the prototypical response seen in young adults.

### 2.7. Similarities of activations during memory encoding (FADE-SAME)

In addition to evaluating the classic FADE score in a large cohort, we further developed the FADE-SAME score as a more comprehensive version of the FADE score, which was motivated based on the following considerations:

1. Older adults do not only deviate in encoding-related fMRI activity from young adults by reduced activations in voxels with a positive effect (*J*_+_), but also by reduced deactivations in voxels with a negative effect (*J*_−_) (see Figure 2; also see Maillet & Rajah, 2014).
2. The normalized activation loss, relative to young subjects, in one voxel is

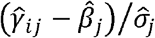

and the normalized deactivation loss, relative to young subjects, in one voxel is

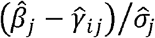

where 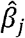 is the average contrast estimate in young subjects, 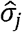 is the standard deviation of young subjects on this contrast at the -*j*th voxel, and 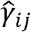 is the contrast estimate of the *i*-th older subject at the *j*-th voxel (see Figure 1).
3. The FADE-SAME score is obtained by averaging within the sets of voxels with positive and negative effect, respectively, and adding up the two components

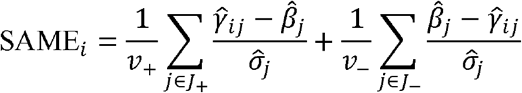

where *ν* _+_ and *ν* _−_ are the numbers of voxels in *J*_+_ and *J*_−_, respectively (see Figure 1B).
4. As becomes evident in this equation, the FADE-SAME score includes a correction for the standard deviation of the parameter estimates at any given voxel in the baseline cohort of young subjects. Thereby, voxels showing prototypical activation across the baseline cohort are weighted more strongly than those activating less robustly.

The FADE-SAME score allows for a number of interpretations (see Appendix A). Most generally, a higher FADE-SAME score indicates higher similarity of an older adult’s brain responses with the activation and deactivation patterns seen in young subjects.

### 2.8. Extraction of FADE-classic and FADE-SAME scores

After single-subject model estimation, FADE-classic and FADE-SAME scores were calculated from t-values (FADE-classic; see Section 2.6) or estimated regression coefficients (FADE-SAME; see Section 2.7). For both the FADE-classic and the FADE-SAME score to be suitable as biomarkers for cognitive aging, it is important to assess to what extent these scores actually reflect age-related activation deviations rather than age-independent individual differences. To explore this potential caveat further, we computed both scores also for the young study participants. In order to avoid circularity issues when calculating scores for a given young subject – whose data were also used to generate reference maps –, the entire cohort was split into two cross-validation (CV) groups.

These CV groups were created by randomly splitting each cohort of subjects (young, older) and then testing whether the two groups significantly differ regarding mean age, gender ratio and scanner ratio. This procedure was repeated until the p-value for all three tests was larger than 0.5 and the final partition was reported for each cohort (see Supplementary Table S3). The resulting CV groups did not differ significantly with respect to (i) their mean age, (ii) the number of male versus female subjects and (iii) the MRI scanner on which participants were investigated (Verio vs. Skyra). Then, scores of any given (young or older) participant in CV group 2 were calculated based on reference maps generated from all young subjects in CV group 1, and vice versa. For completeness, these calculations were also performed for the middle-aged subjects in our cohort (Soch et al., 2021), who were treated as a separate group and are reported in the supplement (see Supplementary Figure S2).

Using this procedure, FADE-classic and FADE-SAME scores were computed on the novelty contrast (novelty – master; contrast vector: *c* = [+ 1,0, − 1]^T^) and on the memory contrast (arcsine-transformed PM; contrast vector: *c* = [0,1,0]^T^), leading to 217 values (number of subjects) for each of the four scores (novelty vs. memory x FADE-classic vs. FADE-SAME) in total.

### 2.9. Statistical evaluation of FADE-classic and FADE-SAME scores

To investigate the robustness and utility of the two scores, the values calculated using the methods described above were subjected to a number of statistical evaluations:

- We first computed between-subject ANOVAs for all scores to test for potential effects of age group, scanner or gender.
- Next, mixed ANOVAs were computed for all scores to test for interactions of the within-subject factor score (FADE-classic vs. FADE-SAME) and the between-subject factor age group (young vs. older).
- To assess relationships between the classic FADE score or the FADE-SAME score and other variables associated with age-related memory decline, we computed correlations with age, memory performance and hippocampal volume within age groups.
  - As an estimate of memory performance, we calculated the area under the ROC curve (A’) from the performance in the memory task performed 70 min after the fMRI experiment (see Appendix B for details).
  - For estimation of hippocampal volumes (V_HC_), individuals’ hippocampal volumes (in mm^3^) were obtained via automatic segmentation with FreeSurfer (Fischl, 2012) and the module for the segmentation of hippocampal subfields and amygdala nuclei (Iglesias et al., 2015; Saygin et al., 2017), which is robust across age groups and MRI scanners (Quattrini et al., 2020) (see Appendix C for details).
- We additionally performed two-sample t-tests between our cohort of young subjects and an independent replication cohort of young subjects (see Section 2.10) in order to assess stability of the scores for young subjects across studies.
- Finally, we calculated correlation coefficients for the scores of older subjects, computed using the young subjects of the main experiment versus the replication subjects as reference, in order to assess stability of the scores for older subjects.

### 2.10. Replication with an independent baseline cohort

The paradigm employed in the present study had previously been used in a cohort of young adults (Assmann et al., 2020; hence termed *yFADE*) consisting of 117 young subjects (60 male, 57 female, age range 19-33, mean age 24.37 ± 2.60 years; see Supplementary Table S1). In the present study, we used those separate young subjects for stability analyses, i.e. (i) to assess whether FADE-classic and FADE-SAME scores are comparable when calculated for young subjects from different cohorts and (ii) to assess whether the two FADE scores are comparable when calculated for older subjects using reference maps from different sets of young subjects.

## 3. Results

### 3.1. Age-related differences in the human memory network could be replicated

Using two-sample t-tests, we compared the age-related activation differences during novelty processing (novel vs. master images) and successful encoding (parametric modulator of the novelty regressor with encoding success). Replicating previous studies (Maillet & Rajah, 2014), we found older participants to exhibit lower activation of inferior and medial temporal structures, particularly of the parahippocampal cortex, but relatively reduced deactivations in midline structures of the DMN during both novelty processing and successful encoding (see Figure 2).

### 3.2. FADE scores are modulated by age, but neither gender nor MRI scanner

We first computed 2×2×2 ANOVAs to assess how the different FADE scores of the 217 subjects in our sample were influenced by (i) the two different age groups (young or older), (ii) gender (male or female) and (iii) the MRI scanners in which they were investigated (Siemens Verio or Skyra; see Section 3.3).

There was a significant main effect of age group for all scores, except for the classic FADE score computed from the novelty contrast, which did not significantly differ between age groups (see Table 2). There were no main effects of scanner, gender, or interactions with them on any of the scores (see Table 2).

**Table 2.** Between-subject ANOVAs for FADE-classic and FADE-SAME scores.

|  | **novelty contrast** |  | **memory contrast** |  |
| --- | --- | --- | --- | --- |
|  | **FADE score** | **SAME score** | **FADE score** | **SAME score** |
| main effect of scanner | F = 0.11, p = 0.742 | F = 0.14, p = 0.707 | F = 0.13, p = 0.723 | F = 1.65, p = 0.201 |
| main effect of gender | F = 0.22, p = 0.636 | F = 1.41, p = 0.236 | F = 0.36, p = 0.550 | F = 2.67, p = 0.104 |
| main effect of age group | F = 0.16, p = 0.686 | F = 16.56,<br>p < 0.001 | F = 81.76,<br>p < 0.001 | F = 135.04,<br>p < 0.001 |
| interaction of scanner and gender | F = 0.05, p = 0.815 | F = 0.00, p = 0.999 | F = 0.06, p = 0.810 | F = 0.20, p = 0.653 |
| interaction of scanner and age group | F = 1.84, p = 0.177 | F = 0.02, p = 0.900 | F = 0.97, p = 0.325 | F = 0.71, p = 0.399 |
| interaction of gender and age group | F = 2.10, p = 0.149 | F = 0.01, p = 0.908 | F = 0.44, p = 0.507 | F = 0.84, p = 0.360 |
| interaction of age group, scanner, and gender | F = 0.40, p = 0.528 | F = 0.05, p = 0.826 | F = 0.00, p = 0.995 | F = 0.03, p = 0.853 |

Results from three-way ANOVAs with scanner, gender and age group as factors for both scores computed from both, novelty and memory contrast. All F-values have one numerator degree of freedom and 209 denominator degrees of freedom.

### 3.3. FADE scores differ in their ability to capture age-related differences

In order to directly compare the modulation of the two scores by age, we additionally computed 2×2 mixed ANOVAs with score (FADE-classic, FADE-SAME) as within-subject factor and age group (young, older) as between-subject factor, separately for the novelty and memory contrasts. There was a significant interaction between score and age for the novelty and memory contrast (see Table 3), supported by larger differences between age groups for the FADE-SAME score. Both scores showed robust age-group-related differences for the memory contrast, and the FADE-SAME score additionally exhibited age-group-related differences for the novelty contrast (see Figure 3).

**Table 3.**
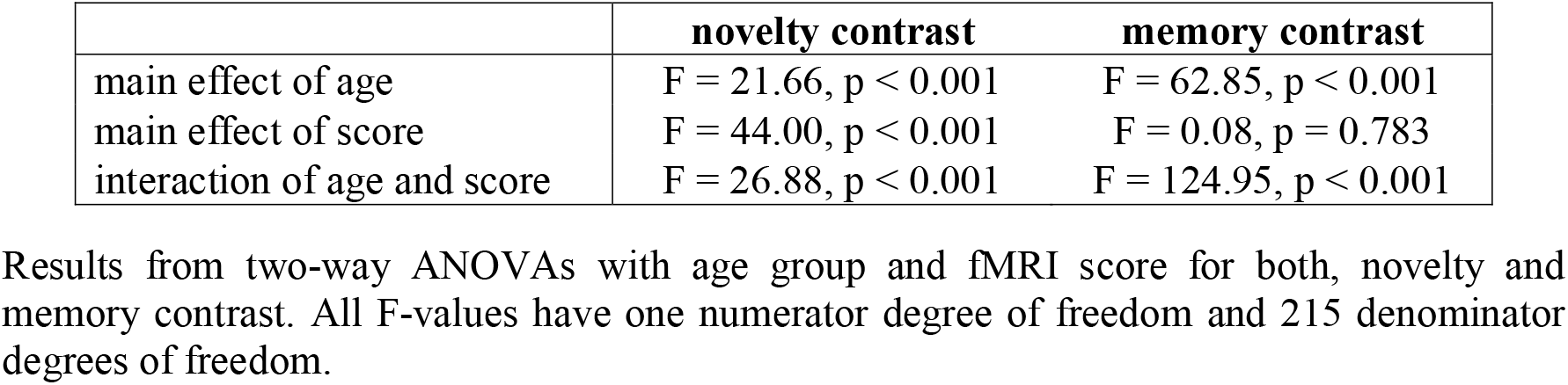
Within-subject ANOVAs for FADE-classic and FADE-SAME scores.

|  | **novelty contrast** | **memory contrast** |
| --- | --- | --- |
| main effect of age | F = 21.66, p < 0.001 | F = 62.85, p < 0.001 |
| main effect of score | F = 44.00, p < 0.001 | F = 0.08, p = 0.783 |
| interaction of age and score | F = 26.88, p < 0.001 | F = 124.95, p < 0.001 |

Results from two-way ANOVAs with age group and fMRI score for both, novelty and memory contrast. All F-values have one numerator degree of freedom and 215 denominator degrees of freedom.

**Figure 3.**
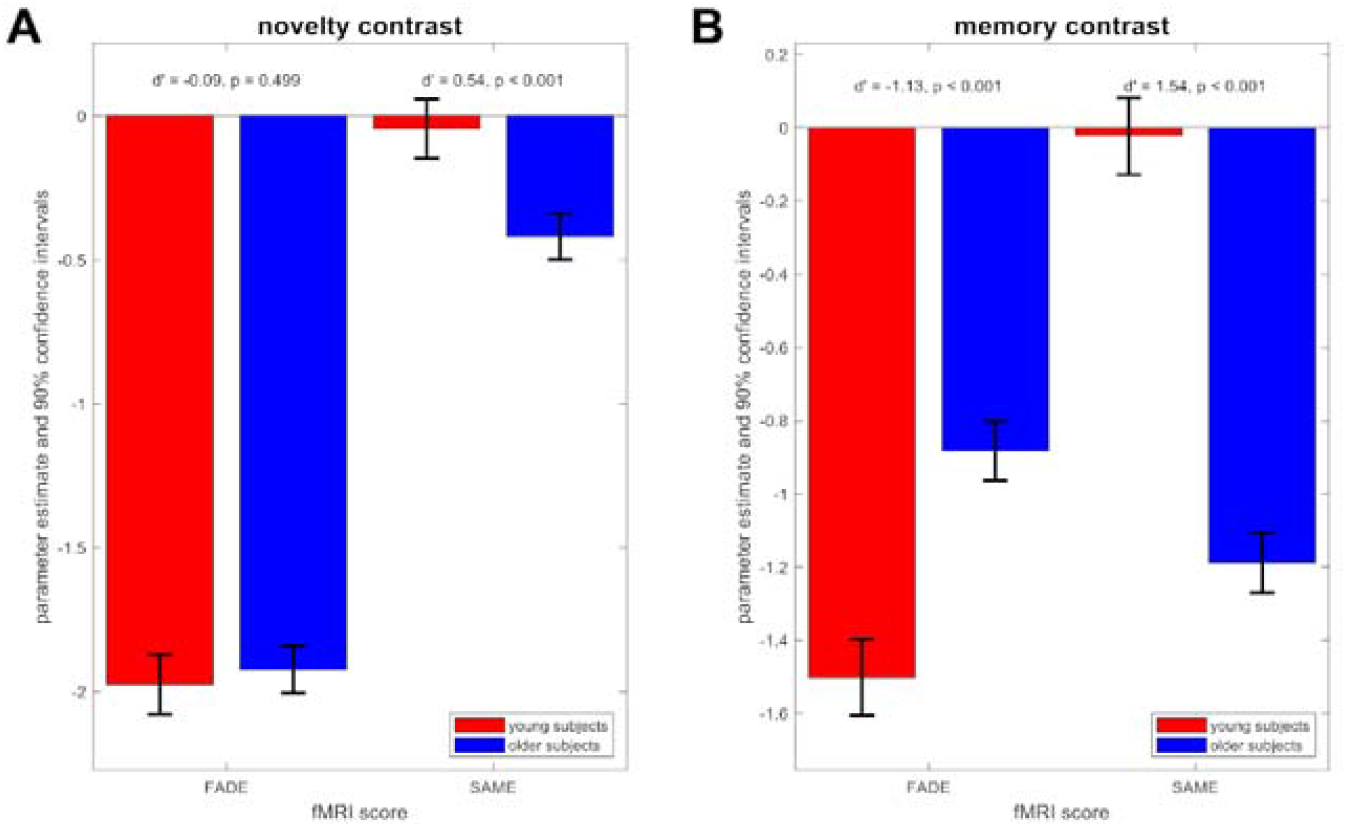
Differences of FADE-classic and FADE-SAME score between age groups. Results from mixed ANOVAs with fMRI score and age group as factors. **(A)** Parameter estimates and 90% confidence intervals for the novelty contrast. The FADE-SAME score shows an age group difference not found for the classic FADE score. **(B)** Parameter estimates and 90% confidence intervals for the memory contrast. Both the classic FADE score and the FADE-SAME score showed pronounced age-related differences.

Due to its construction (see Figure 1), the FADE-SAME score was zero on average for young subjects – because their activation patterns were by definition distributed around the reference activities – and negative on average for older subjects – from summing up activation losses and reduced deactivations. Consequently, the FADE-SAME score was not significantly different from zero across the cohort of young subjects, whereas the negative values in the cohort of older subjects indicates a larger deviation of those individuals’ brain responses from the activation pattern in young subjects (see Figure 3).

When performing the same mixed ANOVAs, but this time comparing older subjects with middle-aged subjects (age range: 51-59 years) instead of young subjects, we found no significant differences between older and middle-aged subjects for any of the scores (see Supplementary Table S4 and Supplementary Figure S2).

### 3.4. FADE scores correlate with other indices of cognitive aging

As described above, the classic FADE score for the memory contrast as well as both FADE-SAME scores differed significantly between age groups. Consequently, these scores were also correlated with age as a continuous variable (FADE-SAME based on memory contrast: r = - 0.63, p < 0.001). However, when controlling for age group, namely calculating separate correlation coefficients for young and older subjects, those correlations were either very low (FADE scores based on memory contrast within older subjects) or not significant (all other scores; see Figure 4, 1^st^ row). Moreover, there were significant correlations with memory performance, as measured by area under the curve (AUC), for the FADE-SAME score and for the classic FADE score computed from the memory contrast (see Figure 4, 2^nd^ row). For the novelty contrast, we observed a significant correlation of the FADE-SAME score, but not of the classic FADE score, with memory performance in older subjects (see Figure 4, 2^nd^ row). No significant correlations with hippocampal volume could be observed for any of the scores (all p > 0.05; see Figure 4, 3^rd^ and 4^th^ row), and we also did not observe robust correlations with the volumes of hippocampal subfields (see Supplementary Table S5).

**Figure 4.**
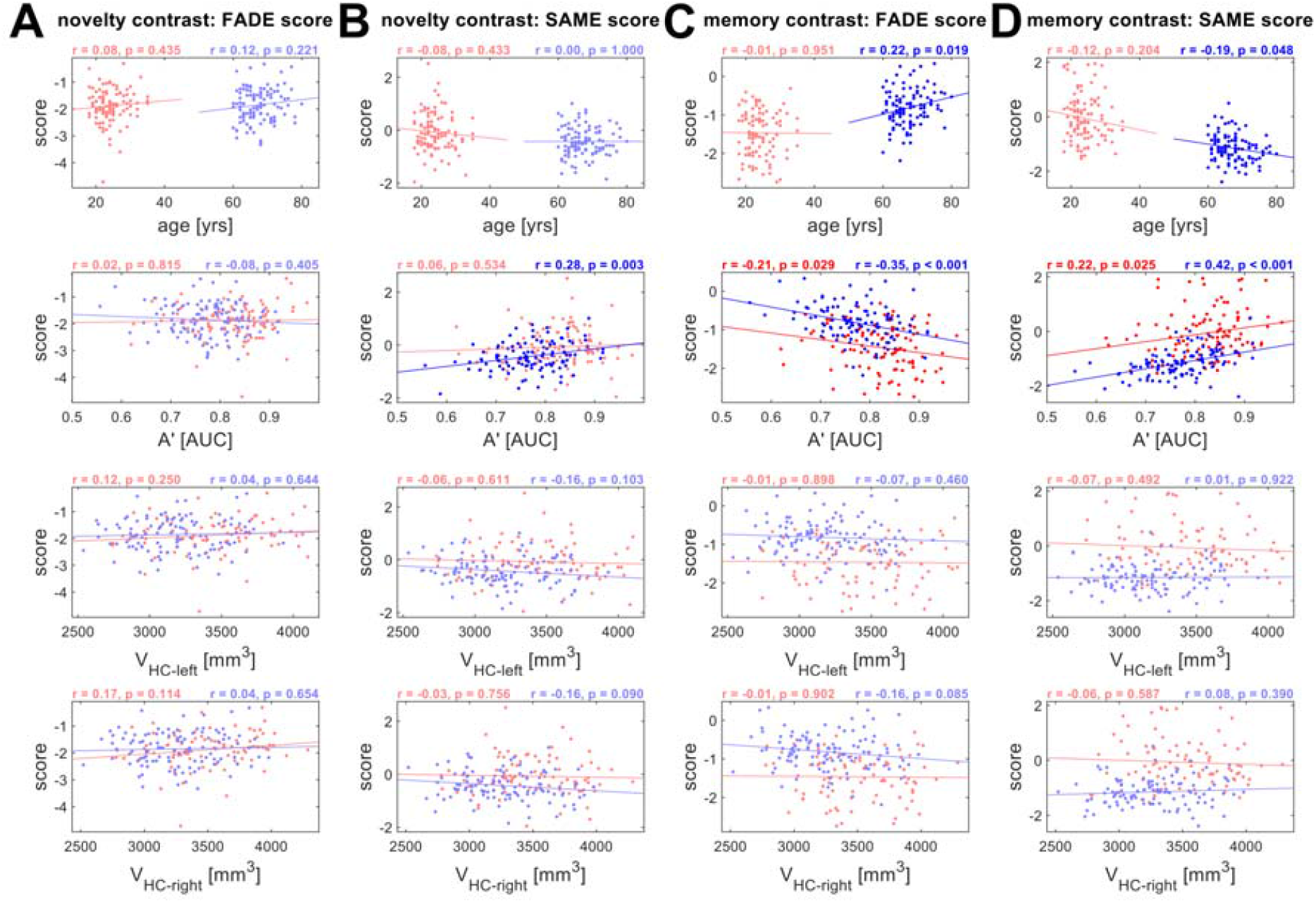
Correlations with independent variables, separated by age group. Results from correlation analyses of FADE-classic and FADE-SAME scores with age, memory performance (A’) and hippocampal volumes (V_HC_). Correlations are reported separately for **(A)** the classic FADE score computed from the novelty contrast, **(B)** the FADE-SAME score computed from the novelty contrast, **(C)** the classic FADE score computed from the memory contrast and **(D)** the FADE-SAME score computed from the memory contrast. Young subjects are depicted in red, and older subjects are depicted in blue. Significant correlation coefficients are highlighted.

To explore the relationship between FADE scores and chronological age further, we plotted the scores as a continuous function of age in years, highlighting that the age dependence of the scores reflects a group effect rather than a continuous relationship with age (see Supplementary Figure S5). In analogy to the correlational analyses depicted in Figure 4, we also report correlations between FADE scores and indices of cognitive aging in the middle-aged subjects (N = 42) and in the young subjects (N = 117) from our replication cohort (see Supplementary Figure S6).

### 3.5. The FADE-SAME score is stable across different cohorts of young subjects

In order to test stability of FADE-classic and FADE-SAME score for young adults, we compared scores obtained from the 106 young subjects in our study sample (see Section 2.1) with scores obtained from the 117 young subjects in the replication cohort (see Section 2.10). Both sets of scores were obtained in a cross-validated fashion, such that all scores were computed using reference maps obtained from independent subjects, but from the same cohort (see Section 2.8).

FADE-SAME scores were close to zero on average by definition (as explained in Section 3.3) and did not differ significantly between original and replication subjects (see Figure 5B/D), whereas classic FADE scores showed significant group differences with small to medium effect sizes for novelty and memory contrast (see Figure 5A/C). Note that both cohorts were comparable regarding age range, mean age, and ratio of male to female participants (see Table 1 and Supplementary Table S1).

**Figure 5.**
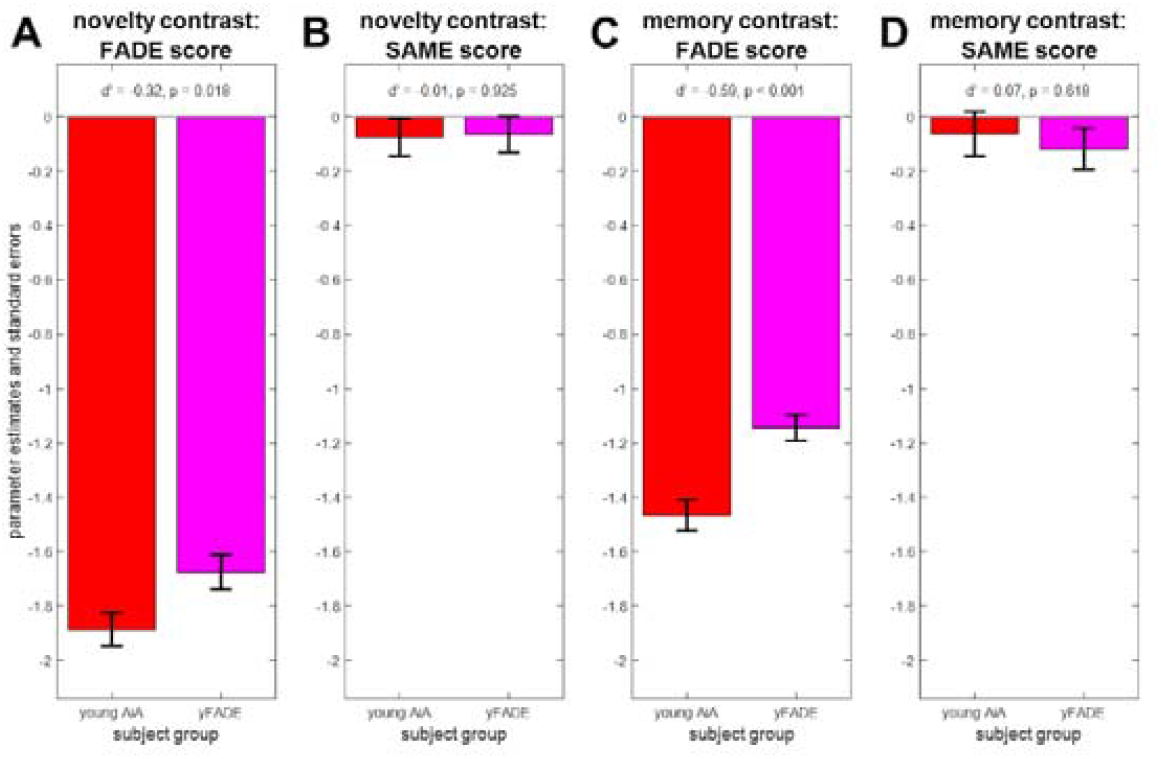
Stability of the FADE scores for young subjects from different studies. Comparison of original young subjects (young AiA, red) and replication young subjects (yFADE, magenta). **(A)** Classic FADE score computed from the novelty contrast. **(B)** FADE-SAME score computed from the novelty contrast. **(C)** Classic FADE score computed from the memory contrast. **(D)** FADE-SAME score computed from the memory contrast. There are no group differences for the FADE-SAME score (B, D), but significant differences between original and replication subjects for the classic FADE score (A, C).

### 3.6. FADE scores are stable for older subjects when using different reference samples

For future use of the classic FADE or FADE-SAME scores in the investigation of older adults and clinical populations, it is important to assess their generalizability, which is, among other factors, determined by their independence from the underlying reference sample. Therefore, we computed both scores for the 111 older subjects in our sample using reference maps (see Figure 1A) obtained either from the young subjects of the main study sample or obtained from the young subjects of the replication cohort. We then calculated Pearson’s correlation coefficients between the scores calculated with the two different baseline samples.

We found all scores (FADE-classic vs. FADE-SAME x novelty vs. memory contrast) to be highly correlated with the respective scores calculated based on the yFADE sample as reference (all r > 0.96, all p < 0.001; see Figure 6), indicating their robustness with respect to different reference samples.

**Figure 6.**
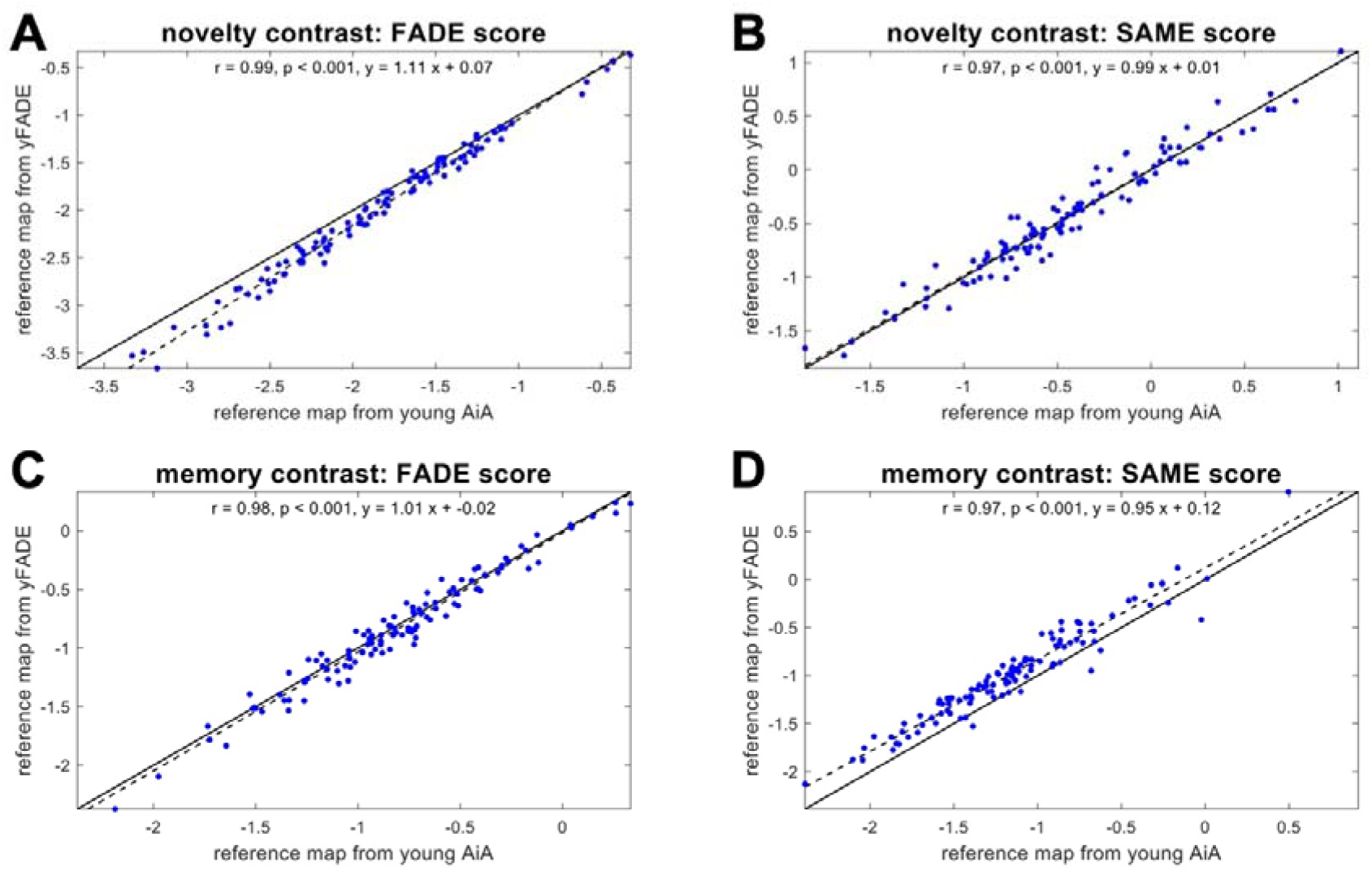
Stability of the FADE scores for older subjects as a function of reference sample. Comparison of scores computed for older subjects (older AiA), using reference maps obtained from either original young subjects (young AiA) or replication young subjects (yFADE). In all panels, the solid black line is the identity function, and the dashed black line represents the regression line. **(A)** Classic FADE score computed based on the novelty contrast. **(B)** FADE-SAME score computed based on the novelty contrast. **(C)** Classic FADE score computed based on the memory contrast. **(D)** FADE-SAME score computed based on the (parametric) memory contrast. There are highly significant correlations for both scores and both contrasts.

In additional analyses, we investigated the correlation between scores computed using reference maps obtained from either all young subjects of one cohort (contrary to the cross-validation scheme used here) or just half of those subjects (roughly equivalent to the cross-validation scheme used here), finding similarly high correlations (see Supplementary Figure S7).

## 4. Discussion

In the study reported here, we have tested the utility of single-value scores of memory-related fMRI activation patterns as potential biomarkers for neurocognitive aging. To this end, we have developed the FADE-SAME score as an enhanced version of the classic FADE score (Düzel et al., 2011), thereby accounting for both individual differences in the baseline sample and the simultaneous presence of activations and deactivations. We then evaluated the two scores (FADE-classic, FADE-SAME), calculated from two different contrasts (novelty processing, subsequent memory) with respect to interpretability, correlation with age and other proxies of age-related memory decline (memory performance, hippocampal volumes) as well as their stability as a function of different reference samples.

### 4.1. Different FADE scores as biomarkers of the aging memory system

Based on the initial work introducing the FADE score as an efficient, reductionist measure of age-related differences of the human MTL memory system (Düzel et al., 2011), we aimed to develop the FADE-SAME score as a more comprehensive measure of age-related differences. To this end, the FADE-SAME score takes into account variability of fMRI activity patterns across the cohort of young subjects used as a reference, and it incorporates differences in encoding-related brain responses in a more holistic way by considering differences in both activations and deactivations (Maillet & Rajah, 2014). Furthermore, we aimed to make the FADE-SAME score more interpretable by defining zero as a fixed value for normalcy, signifying the mean activation pattern of the baseline cohort of young adults.

These theoretical advantages come at the cost of being potentially more dependent on the baseline dataset. Specifically, computing a classic FADE score only requires a set of voxels showing a positive effect in a reference sample of young, healthy subjects. In contrast, computing a FADE-SAME score additionally requires average parameter estimates (i.e., beta values) from the reference set and their standard deviations. This could be a disadvantage of the FADE-SAME relative to the classic FADE score, as, compared to sets of significant voxels, estimated beta values may be more strongly dependent on nuisance variables like different MRI scanners, scanning and preprocessing parameters, or population effects of the chosen baseline sample. All of these factors could – in theory – limit the applicability of FADE-SAME scores for older adults based on activation templates obtained from another study.

Therefore, we aimed to assess the robustness of the different FADE scores with respect to different baseline samples. We calculated scores based on our current study sample and a previously described sample (Assmann et al., 2020) that was demographically comparable, but investigated with slightly shorter trial timings as well as different scanning and preprocessing parameters (Soch et al., 2021). Notably, we observed uniformly strong correlations across the FADE scores based on the two different baseline samples, suggesting that, at least in the case of the present datasets, the aforementioned dependency of the FADE-SAME score on the baseline sample may be negligible in practice (see Figure 6).

For the FADE-SAME score to be employed as a potential biomarker, it is important that it does not merely bear advantages at the theoretical level, but works robustly in empirical investigations. Our validation analyses have indeed revealed highly encouraging empirical evidence regarding the practical utility of the FADE-SAME score. First, the FADE-SAME score yielded a highly robust differentiation between the age groups of young and older subjects, particularly for the memory contrast but also for the novelty contrast (see Figure 3 and discussion below). Second, when controlling for age group, the FADE-SAME score showed significant correlations with behavioral memory performance, not only when computed from the subsequent memory contrast, but also when computed from the novelty contrast (see Figure 4). The latter was not the case for the classic FADE score. Last but not least, when computing FADE scores for the young subjects of our original cohort and the replication cohort, the FADE-SAME scores were associated high stability across subjects, yielding comparable values across the two samples (see Figure 5). This is particularly noteworthy when considering its computationally higher dependence on the reference sample. We suggest that this robustness with respect to the reference sample may be most readily explained by the fact that spurious activations at group level attributable to atypical individual activation patterns (M. B. Miller et al., 2002) are weighted less strongly when accounting for the standard deviation at each voxel.

### 4.2. Novelty and subsequent memory contrasts as basis for the FADE scores

While in the original study by Düzel and colleagues (Düzel et al., 2011), the FADE score was based on neural correlates of successful memory encoding, namely, the DM effect, there is considerable neuroanatomical overlap between the DM effect and the *novelty effect*, which is obtained by comparing novel items to previously familiarized items (Soch et al., 2021). In a previous analysis of memory-related brain activity patterns in young and older adults, we showed that the novelty processing and successful memory encoding engaged largely overlapping networks in the human brain (Soch et al., 2021, Fig. 6A). The analyses reported here have revealed that the same holds true for the age-related activation differences with respect to these contrasts (see Figure 2).

The novelty effect can be computed independently of successful encoding, which may be advantageous in memory-impaired individuals, who have an insufficiently low number of later remembered items for the calculation of a DM effect. However, when computing the FADE-classic and FADE-SAME scores, we found that the FADE scores computed from the subsequent memory contrast showed substantially more robust age differences than those obtained from the novelty contrast. In fact, the classic FADE score computed from the novelty contrast did not discriminate significantly between young and older participants (see Table 3 and Figure 3). This observation was somewhat unexpected, as novelty-related hippocampal activation has already been negatively associated with Tau protein concentrations in the CSF of older adults (Düzel et al., 2018). On the other hand, the FADE scores likely constitute more comprehensive indices of neurocognitive aging than isolated hippocampal activation differences (see Section 3.3). Moreover, a previous extensive longitudinal study revealed that the hippocampus shows either hypo- or hyperactivation during encoding (i.e., encoding vs. control task) in a face-name association task, depending on whether participants age healthily or pathologically, respectively (Nyberg et al., 2019). As our study is cross-sectional, it is conceivable that effects of age-related hypo- and hyperactivation were mixed in our sample, potentially cancelling each other out.

Two notable exceptions to the overall strong overlap of the age-related activation differences of novelty and subsequent memory contrasts are the right dorsolateral prefrontal cortex (dlPFC) and the pregenual anterior cingulate cortex (pgACC). The right dlPFC shows a positive effect of age on novelty (i.e., higher activations for older subjects), but a negative effect of age on memory (i.e. higher activations for young subjects), while the pgACC shows a negative effect of age on novelty, but a positive effect of age on memory (see Figure 2). However, these findings do not seem to contradict the general rule that a region characterized by activations in young subjects shows lower activity in older subjects and that a region characterized by deactivations in young subjects shows reduced deactivations or even absolute activations (relative to baseline) in older subjects (see Figure 2).

### 4.3. The utility of fMRI-based scores as potential biomarkers

Since the first description of the FADE score (Düzel et al., 2011), relatively few studies have used fMRI correlates of memory processes as indices of cognitive aging at the individual level. One study revealed a relationship of dedifferentiation of stimulus-specific processing in the lateral occipital cortex and parahippocampal place area and memory performance (Koen et al., 2019), but in that study, age and memory performance were independently associated with dedifferentiation (for a further discussion, see Koen and Rugg, 2019). Recently, recollection-related fMRI activation of the hippocampus during retrieval has been associated with both memory performance and longitudinal preservation of memory performance in older adults (Hou et al., 2020). While this approach will likely yield similar results to our whole-brain approach with encoding-related activation patterns, it may be limited in subjects with very poor memory performance, like individuals with subjective cognitive decline or mild cognitive impairment, especially considering that an associative word-pair learning task was used. Furthermore, when focusing on the hippocampus, the task-dependency of hippocampal activations must be taken into account. Prior research showed an inverse relationship between CSF tau as a measure of neurodegeneration and increased activation signaling during novelty processing (Düzel et al., 2018). At the same time, the hippocampus showed an – potentially compensatory – over-recruitment and thus more pronounced deviation from the prototypical activation patterns during encoding compared to a low-level baseline (Bookheimer et al., 2000), or during pattern separation (Bakker et al., 2012; Berron et al., 2019). This issue underscores a considerable advantage of whole-brain computed scores like the ones presented here. The exact mechanisms of cognitive reserve in old age are not yet fully understood, but suggest a multitude of factors such as cellular, neurochemical, as well as gray- and white matter integrity, but also functional activation differences (Cabeza et al., 2005; Nyberg et al., 2012). Several fMRI studies point to widespread systems-level activation differences in association with preserved (i.e., little or no differences in comparison to young adults) cognitive and memory performance in particular (Anthony & Lin, 2017; Colangeli et al., 2016; Vidal-Piñeiro et al., 2018). It was beyond the scope of our present work to identify the factors that augment cognitive reserve and/or contribute to brain maintenance (Cabeza et al., 2018), but our exploratory analyses in middle-aged participants suggest a role educational level and established semantic knowledge (see Supplementary Results and Discussion). Research is underway to further characterize the relationship between the scores described here and a more comprehensive neurocognitive profile in older adults (Richter et al., ongoing study).

Irrespective of the specific statistic employed, the current practice for establishing memory-related fMRI activations as a potential biomarker is to compute a summary statistic from an fMRI contrast (e.g., hippocampal activation or a FADE-score-type statistic). The usual aim is that this statistic is linearly related to some clinically relevant variable such as age, memory performance, or grey matter density (see Figure 7A). More recently, partial least squares (PLS)-based decomposition of memory-related fMRI activations has been employed as a whole-brain multivariate approach to identify indices of pathological aging and dementia risk (Rabipour et al., 2020; Salami et al., 2012). In the ongoing search for a memory-related fMRI biomarker with the potential to make predictions at the single-subject level, future studies will be needed to directly compare multivariate and reductionist approaches.

**Figure 7.**
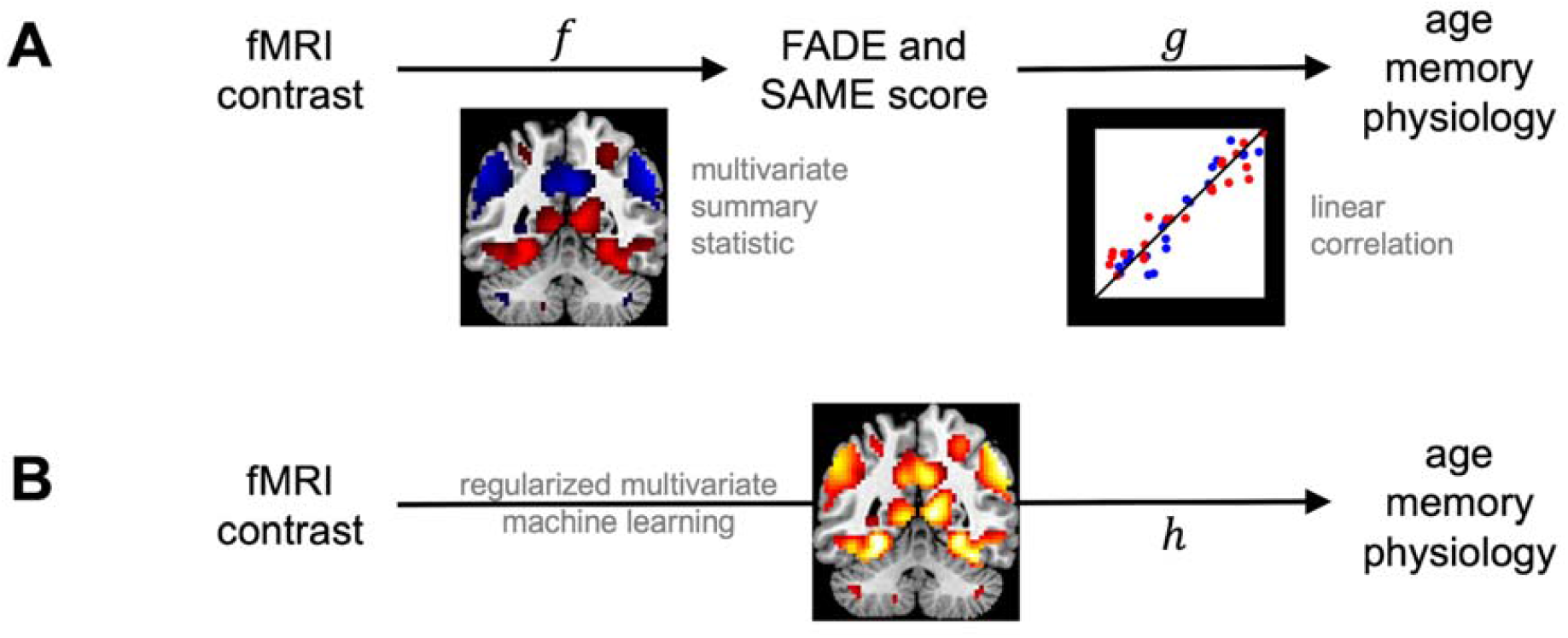
Employing fMRI contrasts to predict human phenotypes. **(A)** Current approach to predicting phenotype from fMRI. A function is calculated from a voxel-wise fMRI contrast map and it is tested whether there is a linear mapping from the outcome of this function to variables of interest. **(B)** Envisaged approach to predicting phenotype from fMRI. The non-linear mapping from voxel-wise fMRI contrast to human phenotype is directly estimated using an advanced machine learning technique.

**Figure 8.**
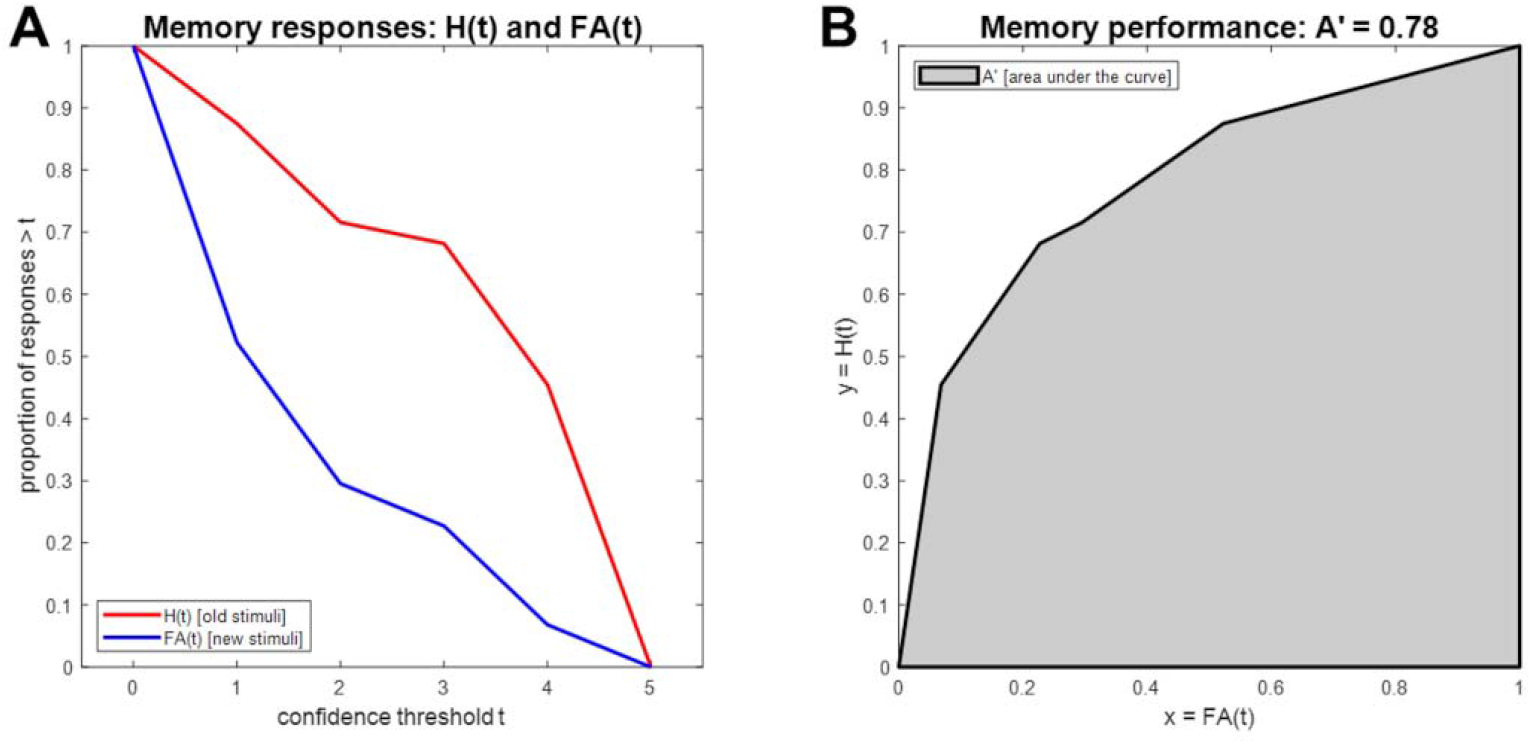
Calculation of A’ as a measure of memory performance. **(A)** Hit rate (H) and false alarm (FA) rate are calculated from raw memory responses as a function of confidence threshold. **(B)** Memory performance is quantified as the area under the curve (AUC) when plotting hit rate against false alarm rate.

In this context, a potentially more sensitive approach might be to directly predict the variable of interest from the voxel-wise fMRI contrast using some regularized multivariate machine learning method (see Figure 7B). Such a method (i) will be based on machine learning, because the precise mapping function is not known *a priori* and (ii) will use regularization, because the number of features (i.e., voxels) is much larger than the number of observations (i.e., patients or healthy older adults) used to train the mapping. For example, one could apply support vector classification (SVC) to decode age group or disease state from estimated fMRI activity; or support vector regression (SVR) to find a non-linear mapping between estimated fMRI activity and observed memory performance, conditional on age. Such an approach may also be helpful in integrating current concepts of successful cognitive aging. While *brain maintenance* is reflected by higher similarity of structure and function with young adults, different forms of *compensation* are rather associated with age-related deviations of brain activity from prototypical patterns found in the young, which are associated with preserved cognitive performance (Cabeza et al., 2018). Non-linear multivariate approaches might in the future allow to consider both mechanisms simultaneously and thereby improve classification and prediction cognitive ability in older adults.

### 4.4. Clinical implications and directions for future research

While investigations of age-related differences in neural correlates of successful memory at the whole-brain level have provided important contributions to our understanding of the neurocognitive changes associated with advanced age (for reviews, see Maillet & Rajah, 2014 and Wang et al., 2010), such single-value biomarkers as presented here may be better suited for clinical applications. The scores can be computed for individual older subjects and compared to a baseline derived from a young reference sample, similarly to classical neuropsychological measures. An important advantage of functional neuroimaging biomarkers over neuropsychological measures is their higher sensitivity for covert age-related changes in episodic memory and potentially additional cognitive domains as, even in the absence of both, differences in behavioral memory performance and obvious structural brain changes, age-related alterations of functional memory networks may be observable (e.g., Cabeza et al., 2002).

In the present study, we restricted our analyses to a neurologically healthy population, and the observed individual differences in the FADE scores therefore most likely reflect physiological inter-individual variability in age-related alterations of the MTL memory system and associated brain networks. In clinical research, the utility of a biomarker depends on its ability to discriminate (i) between healthy controls and affected individuals, or (ii) between different pathophysiological underpinnings of an observed clinical entity. While we were able to show that the evaluated highly reductionist and easy-to-use scores reliably detect correlates of age-related differences in human explicit memory networks, they will yet need to prove their suitability to discriminate, for example, between cognitively impaired individuals with and without underlying Alzheimer’s disease pathology (Jessen et al., 2018).

An important challenge in the evaluation of fMRI-based potential biomarkers for neurocognitive aging is the unclear relationship between hippocampal structure and fMRI-based indices of memory network (dys-)function. Somewhat unexpectedly, none of the scores investigated in the present study correlated robustly with hippocampal volumes. However, this apparent discrepancy is not unique to the present study. In a previous study using the same experimental paradigm (Düzel et al., 2018) in both healthy older adults and individuals with Alzheimer’s disease risk states, CSF tau levels correlated with both hippocampal novelty responses and hippocampal volumes, but these correlations were independent. This observation is in good agreement with earlier reports that memory-related fMRI activations together with ApoE genotype could identify individuals at risk for cognitive decline more accurately than hippocampal volume measures (Woodard et al., 2010). In a combined EEG and structural MRI study, hippocampal volume reductions in older adults were shown to correlate with reduced grey matter density in midline and limbic brain structures, whereas hippocampal diffusivity correlated with more regional grey matter loss in perirhinal and parahippocampal cortices (Schiltz et al., 2006). Notably, only diffusivity was also associated with reduced electrophysiological indices of recollection-based memory, suggesting that hippocampal volumes may not be the optimal predictor of cognitive function in *healthy* older adults. We therefore tentatively suggest that hippocampal volume loss in normal aging and pre-clinical dementia may reflect distinct pathophysiological processes that are differentially related to memory function.

A further question of potentially high clinical relevance will be to what extent the two scores investigated in the present study may correlate with different stages of Alzheimer’s pathology. The classic FADE score constitutes a sum score reflecting reduced activations of the MTL memory system, whereas the FADE-SAME score additionally accounts for age-related hyperactivation (or reduced deactivation) of the brain’s midline structures that constitute the DMN (Maillet & Rajah, 2014). In Alzheimer’s disease, deposition of Tau protein aggregates typically starts in the MTL and subsequently spreads to the brain’s midline. Recently, patterns of Tau deposition could be linked to distinct impairment of item memory versus scene memory (Maass et al., 2019). Future research should thus assess to what extent the scores might differentially reflect MTL versus midline pathology in individuals with Alzheimer’s disease.

### 4.5. Limitations

It is important to note that, while the scores reported here are ultimately aimed at assessing age-related *changes* longitudinally, the current study only assessed *cross-sectional differences* between young and older study participants. Whether the scores are also valuable for predicting changes over time will be assessed in a follow-up study with the large sample of a longitudinal multi-center study of healthy older participants and pre-clinical stages of Alzheimer’s disease (Bainbridge et al., 2019; Düzel et al., 2018).

A further limitation concerns the demographic differences between the samples. They differed with respect to medication status (more chronic diseases with advanced age), and educational background (approximately 50% of the older participants, but 94% of the young participants had obtained the 12-year school-leaving exam). Please see our Supplementary Discussion for likely causes and potential implications of the demographic between-group differences.

Despite the aforementioned differences in demographics, the samples of young and older adults were ethnically and culturally rather homogenous, and all participants (with the exception of two young adults) were of European ancestry. This homogeneity may be considered another limitation, as it might call into question the generalizability of our findings to other populations. Culturally mediated differences in brain activity have been most extensively investigated with respect to European or North American versus East Asian populations and have been most prominently found in social cognitive tasks (Han and Ma, 2014). However, brain activity differences can also be found in non-social tasks like mental arithmetic, which may reflect the shaping of brain networks by literacy in different type systems (Tang et al., 2006). While the use of non-verbal scenes may, in our view, reduce the risk of poor generalizability of our present findings, replication in diverse populations is nevertheless warranted, particularly with respect to memorability of specific images (Bainbridge et al., 2019).

### 4.6. Conclusion

We could demonstrate that single-value scores reflecting age-related deviations from prototypical fMRI activations during memory encoding bear the potential to be used as biomarkers of cognitive aging. Moreover, the FADE-SAME score could also differentiate between age groups when computed from the novelty contrast, suggesting its suitability in memory-impaired clinical populations. In the future, single-value scores reflecting fMRI responses may help to identify distinct subtypes of age-related memory decline and pathological alterations of human memory systems.

## Supporting information

Supplementary Results, Figures, and Tables

## 6. Appendix

### A. Interpretations of the FADE-SAME score

When simply spelling out the equation of the FADE-SAME score, it represents the sum of *average normalized activation loss* in voxels with positive effects and *average normalized deactivation loss* in voxels with negative effects (see Section 2.3):

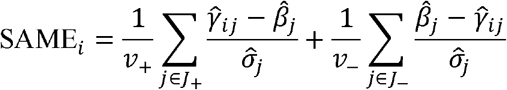

If we focus on just one voxel *j*, then the variance-weighted Euclidean distance of an older adult’s activation from the average young subject in this voxel is

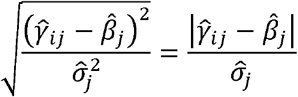

Since we want to obtain directional information (i.e., increased activation/deactivation should benefit while the same amount of decreased activation/deactivation should impair the FADE-SAME score), the activation difference is sign-adjusted for the reference effect

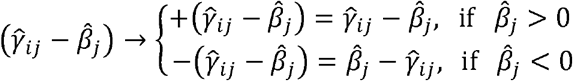

which renders the FADE-SAME score as the *voxel-averaged variance-adjusted directional Euclidean distance* of one subject’s activations from the reference pattern.

Alternatively, one voxel’s term from the sum over voxel sets

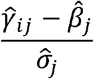

can be seen as an effect size estimate similar to Cohen’s *d* (Cohen, 1988) where the difference of means (or estimated regression coefficients) is divided by the estimated standard deviation. More precisely, the term is equivalent to Glass’ Δ (Glass, 1976)

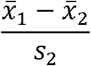

where 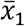 is the estimate from the subject to be assessed (e.g., an older adult), 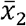 is the average estimate from the control group (i.e., the young adults) and *s*_2_ is the standard deviation calculated from the control group. Taking this into account, the FADE-SAME score is equivalent to the *voxel-averaged directional effect size*.

### B. Computation of memory performance

Let *o*_1_, …, *o*_5_ and *n*_1_, …, *n*_5_ be the numbers of old stimuli and new stimuli, respectively, rated during retrieval as 1 (“definitely new”) to 5 (“definitely old”). Then, hit rates and false alarm (FA) rates as functions of a threshold *t* ∈ {0,1,…,5} are given as the proportions of old stimuli and new stimuli, respectively, rated higher than *t*:

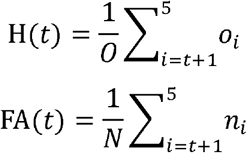

where *O* = *o*_1_, +… +*o*_5_ and *N*= *n*_1_ +… + *n*_5_. Note that H(0) = FA(0) = 1 and H(5) = FA(5) = 0. Consider the hit rate as a function of the FA rate:

*y* = *f*(*x*), such that *y* = H(*t*) and *x* = FA(*t*) for each *t* = 0,1,…,5

Then, the area under the ROC curve is given as the integral of this function from 0 to 1:

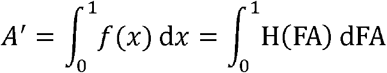

This quantity is referred to as “A-prime” and serves as a measure for memory performance: When the response to each item is random, such that *o*_1_, …, *o*_5_ and *n*_1_, …, *n*_5_ have a uniform distribution, *A* ′ is 0.5, corresponding to pure guessing. When all old items are recognized (*o*_5_ =*O*) and all new items are rejected (*n*_1_ = *N*), *A* ′ is 1, corresponding to perfect performance.

### C. Computation of hippocampal volumes

Hippocampi of individual participants were segmented using FreeSurfer 6.0 and the module for segmentation of hippocampal subfields and amygdalar nuclei^5^, following previous descriptions (Iglesias et al., 2015; Quattrini et al., 2020). In addition to the high-resolution T1-weighted images, high-resolution T2-weighted images acquired perpendicular to the hippocampal axis (see Section 2.3) were processed with the FreeSurfer pipeline, to improve segmentation accuracy (Dounavi et al., 2020). For the purpose of the present study, volumes of the entire hippocampi are reported in the main paper (see Figure 4) and volumes of hippocampal subfields (see Figure S8) are reported in the Supplementary Material (see Table S5).

In the replication cohort (see Supplementary Figure S6), the segmentation with FreeSurfer 6.0 was performed based on T1-weighted MPRAGE images only, as no high-resolution T2-weighted MR images were available in that cohort.

## 7. Statements

## 7.1. Acknowledgments

The authors would like to thank Adriana Barman, Marieke Klein, Kerstin Möhring, Katja Neumann, Ilona Wiedenhöft, and Claus Tempelmann for assistance with MRI data acquisition and Michael Scholz for support with optimization of the fMRI paradigm. We further thank two anonymous reviewers for their thoughtful and constructive comments on a previous version of the manuscript.

## 7.2. Data Availability Statement

Due to data protection regulations, sharing of the entire data set underlying this study in a public repository is not possible. We have however provided GLM beta images as a NeuroVault collection (https://neurovault.org/collections/EPLZNQAD/) for a total of 217 (original cohort) + 42 (middle-aged subjects, see Supplementary Information) + 117 (replication cohort, see Section 2.10) = 376 subjects. Access to de-identified raw data will be provided by the authors upon reasonable request. MATLAB code and instructions to process the data can be found in an accompanying GitHub repository (https://github.com/JoramSoch/FADE_SAME).

## 7.3. Funding and Conflict of Interest declaration

This study was supported by the State of Saxony-Anhalt and the European Union (Research Alliance “Autonomy in Old Age”) and by the Deutsche Forschungsgemeinschaft (SFB 779, TP A08 and SFB 1436, TP A05, to B.H.S. and C.S; DFG RI 2964-1 to A.R.). The funding agencies had no role in the design or analysis of the study. The authors have no conflict of interest, financial or otherwise, to declare.

## Footnotes

1 Please note that we are only considering scores based on task-related fMRI for the current study. While resting-state fMRI-based assessments of cognitive aging have also been evaluated by others (e.g., Sperling, 2011), they follow a very different rationale. One advantage of task-related fMRI is the specificity of task-related activation for concrete cognitive functions. We can differentiate between functional differences related to, for example, long-term memory encoding, novelty detection, working memory, or attention. Thus, it can potentially be used for testing age-related functional differences in a specific cognitive domain (here: episodic memory encoding). Although resting-state fMRI provides a fast and easy approach to assess all fMRI group differences at once, it cannot be used to this end, but rather for assessing general age-related functional differences.

2 The sample also included a smaller subgroup of middle-aged individuals (N = 42; age range: 51-59 years) who were of lesser interest for the current analyses, but whose data are reported in the Supplementary Material for completeness reasons.

3 Colloquially, the memory experiment employed here is also called the “FADE paradigm” due to this score.

4 Note that this is stricter than in the original publication where a significance level of p < 0.001, uncorrected, minimum cluster size k = 6 voxels was applied.

5 URL: https://surfer.nmr.mgh.harvard.edu/fswiki/HippocampalSubfieldsAndNucleiOfAmygdala.

## Notes

https://github.com/JoramSoch/FADE_SAME

### Competing Interest Statement

The authors have declared no competing interest.

### Summary of Updates

Additional revisions as requested by peer reviewers, including analyses controlling for educational background.

