## Supplementary Results, Figures, and Tables for "A comprehensive score reflecting memory-related fMRI activations and deactivations as potential biomarker for neurocognitive aging"

### Supplementary Methods

#### Supplementary sample information

In addition to the 106 young subjects and 111 older subjects (see Section 2.1 in the main paper), our complete study cohort also included 117 young replication subjects (see Section 2.10 in the main paper) and 42 middle-aged subjects (13 male, 29 female, age range 51-59, mean age 55.48 ± 2.57 years) who were part of the sample investigated in an earlier study (Soch et al., 2021). Demographic information of middle-aged and replication subjects is given in Table S1 and medication received by young, middle-aged and older subjects is given in Table S2.

As described in the main paper (see Section 2.8), the subjects from each cohort (young, older, middle-aged, replication) were randomly split into cross-validation groups (see Table S3).

#### Genotyping for apolipoprotein E

Genomic DNA was extracted from blood leukocytes using the GeneMole automated DNA extraction system (GeneMole). Genotyping of the apolipoprotein E (ApoE) haplotypes (ε2, ε3, ε4) was performed using PCR followed by restriction fragment length analysis, as described previously (Addya et al., 1997), with minor modifications. Briefly, the 270 bp fragment containing the single nucleotide polymorphisms rs429358 and rs7412 was amplified using PCR with the primers (ApoE-f: GCA CGG CTG TCC AAG GAG CTG CAG GC; ApoE-r: GGC GCT CGC GGA TGG CGC TGA G; see (Tsukamoto et al., 1993), followed by digestion with Hha I (New England Biolabs) at 37°C overnight. The resulting DNA fragments (ε2: 83+91 bp; ε3: 35+48+91 bp; ε4: 35+48+72 bp) were separated on a 5% agarose gel stained with MidoriGreen and visualized under UV light. To ensure robustness of genotyping, we verified our results using the protocol described by Ossendorf and Prellwitz (2000).

#### Comparison of older subjects by memory performance

To test for potential differences in memory-related brain activations as a function of memory performance, older subjects were divided into four groups (Q1, Q2, Q3, Q4) based on their memory performance (A'), as calculated for correlational analyses (see Appendix B in the main paper), and two of these groups were selected as high-performing (Q4, top quarter) and low-performing (Q1, bottom quarter) older adults.

Then, novelty and memory contrast estimates of subjects from these two groups as well as young subjects were compared using a full factorial design in SPM, with age group (young, Q1, Q4) as a between-subject factor.

**Further analyses of the different FADE scores**

- To test for a potential difference between middle-aged and older subjects regarding biomarkers of neurocognitive aging, we computed mixed ANOVAs with score (FADE-classic, FADE-SAME) as within-subject factor and age group (middle-aged, older) as between-subject factor, similar to the analyses reported in Figure 3 and Table 3.
- To further illustrate the distribution of each score (FADE-classic/FADE-SAME x novelty/memory) as a function of age, we plotted all scores against age in years and calculated smooth mean and smooth variance using a sliding window approach.
- Because the ApoE gene has been previously linked to impaired cognitive function and Alzheimer's disease (Bookheimer et al., 2000), we compared E3 homozygotes (considered the neutral genotype) against E4 carriers (considered the risk allele). To this end, we computed two-way ANOVAs with age group (young, middle-aged, older) and ApoE genotype (E3/E3 vs. E3/E4 or E4/E4) as between-subject factors.
- Because there were no correlations between FADE scores and left or right hippocampus volume (see Figure 4), we also computed correlations of the scores with volumes of hippocampal subfields, as extracted via FreeSurfer (see Appendix C), separated by hemisphere and age group (see Table S5).

**Extraction of FADE scores from the whole versus split sample of young subjects**

Finally, we assessed the dependence of FADE scores on the number of young subjects contributing to the respective reference maps. We have already shown that FADE scores are highly correlated when using independent cohorts of young subjects as reference samples (see Figure 6 in the main paper). Here, we additionally investigated how a smaller or larger reference set of young subjects might impact the FADE scores of older subjects. To this end, the scores of older subjects were extracted using reference maps obtained from either (i) all young subjects from the main cohort (N = 106), (ii) the first half of those subjects (CV group 1, N = 53) or (ii) the second half of those subjects (CV group 2, N = 53). Then, we computed correlations between scores (i) and (ii) as well as (i) and (iii) and calculated the corresponding type 3 intra-class correlation (ICC) coefficients of the means, where the reference samples (i), (ii), (iii) were regarded as different raters of the same outcome variable (i.e. the score).

### Supplementary Results

**Lack of robust differences by memory performance in older subjects**

To investigate if young subject's brain responses are more comparable to older subjects performing high vs. low in the recognition memory task, we calculated a one-way ANOVA of young subjects against high-performing (Q4) and low-performing (Q1) older adults. There was a significant effect of group on novelty and memory contrast estimates in some parts of the human memory network. However, these effects were largely attributable to a difference between young and older subjects rather than a difference between Q1 and Q4 subjects (see Figure S1).

When assessing novelty and memory contrast in voxels with significant differences between young and older subjects (see Figure 2 in the main paper), (i) there were no differences between high- and low-performing older adults in two regions (left (P)HC, pgACC); and (ii) in the other two regions (PreCun, right dlPFC), there was a tendency of activations in high-performing older adults to be more similar to activations in young subjects (see Figure S1). However, when directly comparing Q1 and Q4, there were no activation differences that survived whole-brain correction for multiple comparisons (FWE, $p < 0.05$, $k = 10$).

Note that these voxel-wise analyses do not preclude the possibility of multi-voxel differences in older subjects with high versus low memory performance. In fact, we observed that memory performance correlated with both FADE scores computed from the memory contrast (see main manuscript, Figure 4C and 4D).

**FADE scores do not differ significantly between middle-aged and older subjects**

There was no significant difference between middle-aged and older subjects on any of the two scores computed from either novelty or memory contrasts (see Figure S2). Specifically, in our mixed ANOVA models, there were main effects of score on both contrasts, but no main effects of age and no interactions between age and score (see Table S4).

**FADE scores do not differ significantly as a function of ApoE genotype**

There were no significant differences between any of the four FADE scores of ApoE ε3 homozygotes and ε4 carriers (see Figure S3). Specifically, in our two-way ANOVA models, there were main effects of age group on all scores except the classic FADE score computed from the novelty contrast (cf. Table 2 in the main paper), but no main effect of ApoE genotype and no interaction between ApoE genotype and age group.

**FADE scores are age-independent within age groups**

Visualizing FADE scores as a continuous function of age (see Figure S5) illustrates several results described in detail in the main paper, namely that (i) the FADE-SAME score extracts age-related information from both novelty and memory contrasts, while the classic FADE score showed an age-group difference on the memory contrast only (see Figure 3); (ii) the FADE-SAME score was zero on average for young subjects and negative on average for older subjects (see Figures 3 and 5); (iii) there were differences between age groups for almost all scores (see Table 2), but at maximum low correlations with age within age groups (see Figure 4); and (iv) scores of middle-aged subjects did not differ substantially from scores of older subjects (see Figure S2 and Table S4).

**FADE scores do not correlate with hippocampal subfield volumes**

There were nominally significant correlations between the FADE-SAME score computed from the novelty contrast and left hippocampal tail ($r = -0.25$, $p = 0.008$) as well as right hippocampal tail ($r = -0.24$, $p = 0.010$), both in older subjects (see Table S5). However, these did not survive Bonferroni correction for number of hippocampal subfields (2 x 5 = 10).

**More reference subjects reduce variability of FADE scores**

Our last analysis revealed that FADE scores of older subjects, computed using either the complete sample of young subjects (x; N = 106) or only half of those subjects (y; N = 53), were highly correlated (all $r > 0.98$, all ICC > 0.96, all $p < 0.001$; see Figure S7). However, invariably across contrasts, scores and CV groups, we noted that (i) the intercept of the regression line was close to zero and (ii) its slope was larger than one, typically around 1.2. This signifies that (i) scores are not shifted by increasing or reducing the reference set, but (ii) instead, scores are more variable when computing them based on a smaller reference set.

The explanation for this is that the higher statistical power resulting from a larger reference sample allows one to detect more voxels as being significantly different from zero on a given contrast. Consequently, more voxels – including voxels with smaller effects at group level – contribute to the calculation of the scores, making them smaller overall. We therefore suggest that, when focusing on neural processes underlying age-related memory decline rather than age group comparisons, scores should be calculated based on all available young subjects.

### Supplementary Discussion

**Demographic differences between the experimental cohorts**

While we took great care to ensure that both samples, that is, the young and older adults, were neurologically and psychiatrically healthy and showed no signs of cognitive impairment, we found significant between-group differences with respect to ApoE genotype distribution, medication status, and educational background.

***Differences in ApoE genotype distribution:*** Older, but not young or middle-aged, participants show a lower frequency of the ApoE ε4 allele and a higher frequency of the ε2 allele when compared to a large, ethnically comparable sample from another study (X. Li et al., 2019; see Figure S3A). While the ε4 allele has been robustly associated with increased risk for Alzheimer's disease, the ε2 allele rather convers a protective effect (Z. Li et al., 2020). Beyond dementia risk, ApoE genotype has been associated with longevity, with the ε4 allele being less common in very old age, whereas the ε2 allele is more prevalent in individuals at very advanced age (Deelen et al., 2019). Considering the increased risk of ApoE ε4 carriers to develop Alzheimer's disease and also cardiovascular disease (Marais, 2019), especially from the age of 60 onwards, an underrepresentation of ε4 carriers in a sample of generally very healthy older adults may thus reflect a largely unavoidable selection bias.

***Differences in medication status:*** As expected, medication status differed between age groups (see Table S2), reflecting the higher prevalence of common chronic conditions like hypertension, diabetes, or hypothyroidism with increasing age and the common use of oral contraceptives in young women. Similar to the distribution of ApoE genotypes, we consider these differences largely unavoidable, as they reflect very common factors closely associated with age. Excluding participants taking any medication might, in our view, have led to a selection of an atypically healthy cohort of older participants not representative of the general population. Such a rigid selection may be helpful for certain carefully controlled studies of complex cognitive function in older adults, but less so for the identification of potential neurocognitive biomarkers that reflect the interindividual variability among older adults.

***Differences in educational background:*** While differences in ApoE genotype distribution and medication status likely reflect unavoidable factors closely associated with age (see above), the differences in educational background warrant explanation. Approximately 50% of the older participants, but 94% of the young participants had obtained the 12-year school-leaving exam

(*Abitur*). In our view, this discrepancy is best explained by historically grounded differences in educational systems. Unlike in studies from English-speaking countries, the educational system in Germany, and particularly East Germany where the present study was conducted, has undergone profound changes over the past decades. In the German Democratic Republic (GDR), access to university-level education was tightly regulated by the federal government and depended not only on academic achievement, but also on social and political factors unrelated to individual cognitive capabilities. In the 1970s and 1980s, Abitur was obtained by no more than 12% to 15% of students in the GDR (Köhler et al., 2001). After the German reunification in 1990, school systems in the East German states were restructured and aligned with those in the West German states, and the percentage of students obtaining Abitur increased to approximately 30% (Statistisches Landesamt Sachsen-Anhalt, 2017).

Importantly, despite the on average formally lower educational level in the older participants, it should be noted that they showed higher performance in the MWT-B, a vocabulary-based screening of verbal intelligence (Lehrl, 2005). In our view, it therefore seems safe to assume that both young and older participants had an above-average educational background and average to high cognitive ability.

**The relationship between FADE scores and educational level**

When analyzing FADE scores from subjects with vs. without the Abitur, we found main effects of educational status on the memory contrast, but these effects were largely driven by middle-aged subjects, with no differences in older subjects (see Figure S4B). As discussed above with respect to ApoE genotypes, we cannot exclude that this may result from a selection bias in our cohort of older adults. Alternatively, or more likely additionally, this observation may point to the possibility that the well-known role of educational level as a neurocognitive reserve (Stern, 2012) may be of particular importance in the transition from middle to old age (i.e., between 50 and 60 years).

When correlating FADE scores with MWT-B hit rates, we found no significant correlations in young, older or middle-aged subjects, although trends emerged for correlations with the memory contrast-derived scores in the middle-aged group (see Figure S4A). If these correlations were to prove robust in future replication attempts, this would also be in line with the aforementioned notion of educational level and acquired knowledge as a relevant cognitive reserve in the transition phase form middle to old age.

### Supplementary Tables

**Table S1.** *Demographics for the young replication cohort and middle-aged subjects.*

| | **replication subjects** | **middle-aged subjects** | **statistics** |
|---|---|---|---|
| N | 117 | 42 | – |
| age range | 18-33 yrs | 51-59 yrs | – |
| mean age ± SD | 24.37 ± 2.60 yrs | 55.48 ± 2.57 yrs | $t = -66.81, p < 0.001$ |
| gender ratio | 60/57 m/f | 13/29 m/f | $\chi^2 = 5.14, p = 0.023$ |
| ethnic composition | 117 European | 42 European | – |
| educational status | – | 16/26<br>with / without Abitur | – |
| ApoE genotype | – | 1/6/0/25/8/2<br>E2/E2 / E2/E3 / E2/E4 / E3/E3 / E3/E4 / E4/E4 | – |
| mean MMSE performance ± SD | – | 29.19 ± 0.86<br>(range: 27-30) | – |
| MWT-B hits ± SD | – | 29.59 ± 3.76<br>(1 missing) | – |

Demographic information for young replication subjects (see Section 2.10 in the main paper) and middle-aged subjects (see Supplementary Methods), along with statistics from a two-sample t-test (mean age) and a chi-squared test (gender ratio). Abbreviations: N = sample size, SD = standard deviation, yrs = years, m = male, f = female, MMSE = Mini-Mental State Examination (Creavin et al., 2016; Folstein et al., 1975), MWT-B = multiple choice vocabulary intelligence test ("Mehrfachwahl-Wortschatz-Intelligenztest"; Lehrl, 1999). "Abitur" is the German equivalent of a high school graduation certificate. This table mirrors Table 1 from the main paper.

**Table S2.** *Medication and history of endocrine-related surgery, separated by age group.*

| | young subjects | older subjects | middle-aged subjects |
|---|---|---|---|
| N | 106 | 111 | 42 |
| oral antidiabetics | 0 | 1 | 0 |
| antihypertensives | 2 | 44 | 5 |
| statins | 0 | 4 | 0 |
| thyroid hormone | 1 | 17 | 1 |
| contraceptives and/or hormone replacement medication | 28 | 2 | 5 |
| history of oophorectomy | 0 | 2/2 unilateral/bilateral | 0/1 unilateral/bilateral |
| history of thyroidectomy | 0 | 2 | 0 |

Medication received by young, older and middle-aged, according to self-report in health questionnaire. This table adds information for subjects reported in Tables 1 and S1.

**Table S3.** *Splitting of cohorts into cross-validation groups.*

| Cohort | Variable | Group 1 | Group 2 | Test |
|---|---|---|---|---|
| young subjects | subjects | N = 53 | N = 53 | |
| | mean age | 23.89 ± 3.66 yrs | 24.36 ± 4.34 yrs | t = -0.60, p = 0.547 |
| | gender ratio | 24 : 29 m/f | 23 : 30 m/f | OR = 1.08, p = 1.000 |
| | scanner ratio | 28 : 25 V/S | 30 : 23 V/S | OR = 0.86, p = 0.845 |
| older subjects | subjects | N = 56 | N = 55 | |
| | mean age | 67.34 ± 4.88 yrs | 67.22 ± 4.45 yrs | t = 0.14, p = 0.892 |
| | gender ratio | 22 : 34 m/f | 24 : 31 m/f | OR = 0.84, p = 0.702 |
| | scanner ratio | 33 : 23 V/S | 31 : 24 V/S | OR = 1.11, p = 0.849 |
| middle-aged subjects | subjects | N = 21 | N = 21 | |
| | mean age | 55.29 ± 2.55 yrs | 55.67 ± 2.63 yrs | t = -0.48, p = 0.637 |
| | gender ratio | 6 : 15 m/f | 7 : 14 m/f | OR = 0.80, p = 1.000 |
| | scanner ratio | 9 : 12 V/S | 10 : 11 V/S | OR = 0.82, p = 1.000 |
| replication subjects | subjects | N = 59 | N = 58 | |
| | mean age | 24.27 ± 2.38 yrs | 24.48 ± 2.81 yrs | t = -0.43, p = 0.669 |
| | gender ratio | 29 : 30 m/f | 31 : 27 m/f | OR = 0.84, p = 0.713 |
| | scanner ratio | 59 : 0 V/S | 58 : 0 V/S | n.d. |

For each cohort of subjects, mean age, gender ratio and scanner ratio are given for the two cross-validation groups, along with statistics from a two-sample t-test (mean age) and Fisher's exact tests (gender ratio, scanner ratio). Note that this table also includes the cohorts of middle-aged subjects (see Supplementary Methods) and young replication subjects (see Section 2.10 in the main paper). Abbreviations: yrs = years, m = male, f = female, V = Verio, S = Skyra, OR = odds ratio.

**Table S4.** *Comparison of older and middle-aged subjects.*

| | **novelty contrast** | **memory contrast** |
|---|---|---|
| main effect of age | F = 0.15, p = 0.704 | F = 0.89, p = 0.346 |
| main effect of score | F = 76.55, p < 0.001 | F = 168.05, p < 0.001 |
| interaction of age and score | F = 1.18, p = 0.279 | F = 0.09, p = 0.768 |

Results from two-way ANOVAs with age group and scores for both, novelty and memory contrasts. All F-values have one numerator degree of freedom and 153 denominator degrees of freedom. This table mirrors Table 3 from the main paper.

**Table S5.** *Correlations of scores with hippocampal subfield volumes.*

| **HC** | **Subfield** | **Cohort** | **novelty contrast: FADE score** | **novelty contrast: SAME score** | **memory contrast: FADE score** | **memory contrast: SAME score** |
|---|---|---|---|---|---|---|
| left | Tail | young | r = 0.14, p = 0.159 | r = -0.16, p = 0.101 | r = 0.08, p = 0.418 | r = -0.11, p = 0.279 |
| | | older | r = 0.13, p = 0.158 | r = -0.25, p = 0.008* | r = -0.11, p = 0.258 | r = 0.09, p = 0.356 |
| | Sub | young | r = 0.08, p = 0.396 | r = -0.02, p = 0.827 | r = 0.00, p = 0.961 | r = -0.06, p = 0.515 |
| | | older | r = 0.01, p = 0.894 | r = -0.12, p = 0.212 | r = -0.09, p = 0.347 | r = 0.05, p = 0.611 |
| | CA1 | young | r = 0.05, p = 0.626 | r = -0.01, p = 0.939 | r = -0.01, p = 0.948 | r = -0.07, p = 0.468 |
| | | older | r = 0.06, p = 0.540 | r = -0.14, p = 0.150 | r = -0.01, p = 0.891 | r = -0.05, p = 0.629 |
| | CA3 | young | r = 0.01, p = 0.933 | r = -0.05, p = 0.620 | r = 0.10, p = 0.287 | r = -0.11, p = 0.269 |
| | | older | r = -0.02, p = 0.867 | r = -0.15, p = 0.126 | r = -0.05, p = 0.573 | r = 0.01, p = 0.899 |
| | CA4 | young | r = 0.05, p = 0.625 | r = -0.00, p = 0.988 | r = -0.01, p = 0.913 | r = -0.07, p = 0.483 |
| | | older | r = -0.05, p = 0.586 | r = -0.08, p = 0.390 | r = -0.14, p = 0.155 | r = 0.02, p = 0.845 |
| right | Tail | young | r = 0.10, p = 0.312 | r = -0.04, p = 0.718 | r = 0.02, p = 0.812 | r = -0.07, p = 0.483 |
| | | older | r = 0.10, p = 0.296 | r = -0.24, p = 0.010* | r = -0.16, p = 0.085 | r = 0.11, p = 0.237 |
| | Sub | young | r = 0.17, p = 0.076 | r = -0.05, p = 0.616 | r = -0.07, p = 0.508 | r = -0.01, p = 0.928 |
| | | older | r = -0.08, p = 0.399 | r = -0.01, p = 0.952 | r = -0.18, p = 0.062 | r = 0.05, p = 0.578 |
| | CA1 | young | r = 0.09, p = 0.386 | r = 0.03, p = 0.763 | r = 0.03, p = 0.771 | r = -0.11, p = 0.277 |
| | | older | r = 0.06, p = 0.523 | r = -0.13, p = 0.184 | r = -0.12, p = 0.196 | r = 0.06, p = 0.560 |
| | CA3 | young | r = 0.02, p = 0.824 | r = -0.01, p = 0.953 | r = 0.13, p = 0.170 | r = -0.13, p = 0.194 |
| | | older | r = 0.05, p = 0.602 | r = -0.18, p = 0.059 | r = -0.09, p = 0.325 | r = 0.09, p = 0.332 |
| | CA4 | young | r = 0.02, p = 0.814 | r = 0.04, p = 0.707 | r = -0.00, p = 0.991 | r = -0.07, p = 0.487 |
| | | older | r = -0.01, p = 0.911 | r = -0.10, p = 0.289 | r = -0.16, p = 0.088 | r = 0.08, p = 0.412 |

Results from correlation analyses of FADE-classic and FADE-SAME scores with volumes of hippocampal subfields; *p < 0.05. This table more closely investigates correlations reported in Figure 4 of the main paper.

### Supplementary Figures

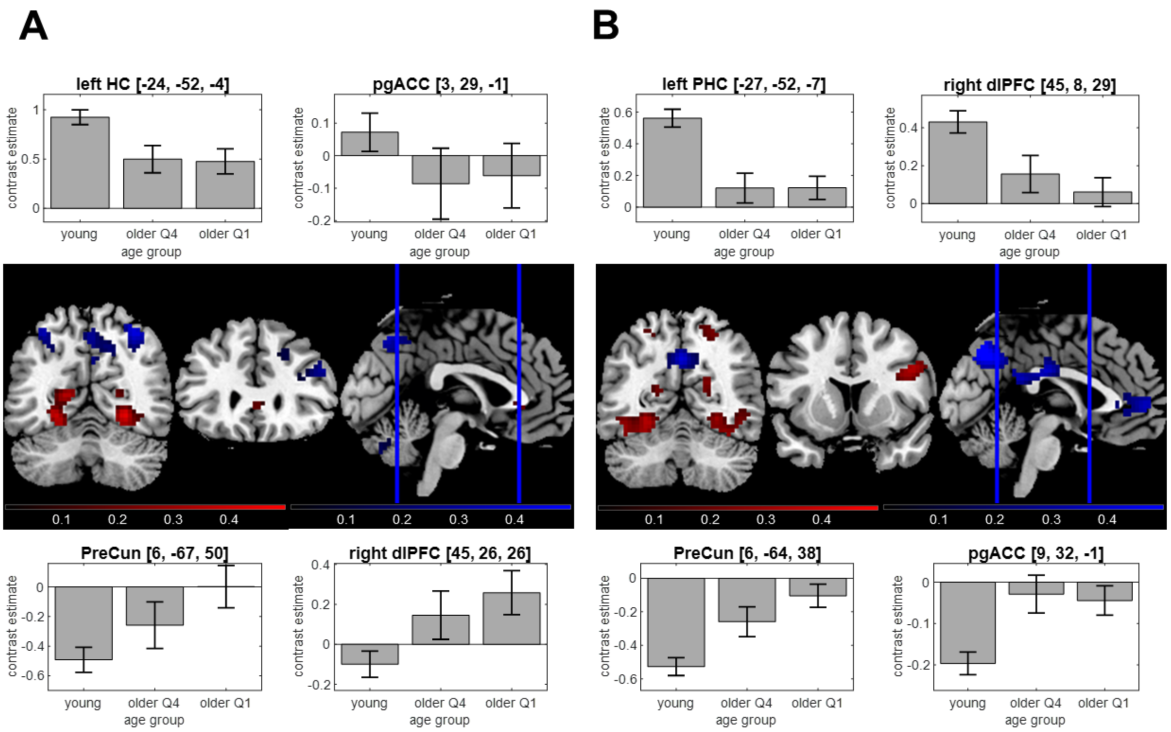


**Figure S1.** *Performance-related differences in the human memory network.* In voxels with significant differences between young and older subjects (see Figure 2), we assessed **(A)** novelty contrast and **(B)** memory contrast for young, high-performing (Q4) and low-performing (Q1) older adults (top and bottom quarter w.r.t. memory performance). Brain sections show significant differences between young and older subjects for activations (red) and deactivations (blue) in young subjects. Bar plots show group-level contrast estimates (gray) and 90% confidence intervals.

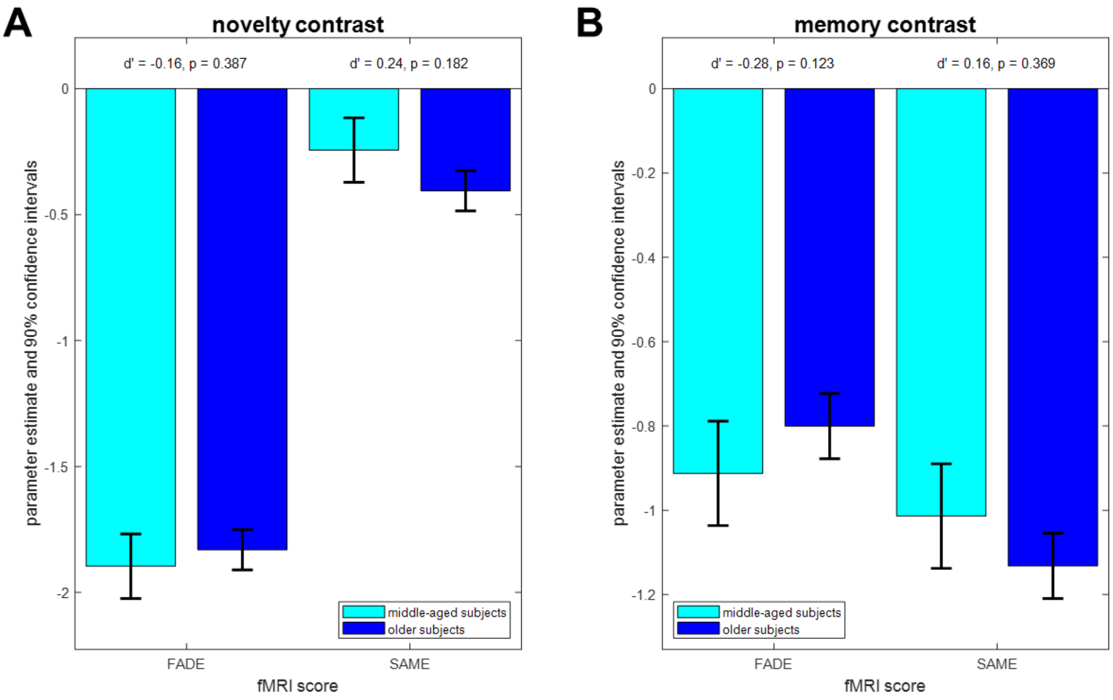


**Figure S2.** *Comparisons of FADE scores from older and middle-aged subjects.* Results of mixed ANOVAs with score as within-subject factor and age group between-subject factor. **(A)** Parameter estimates and 90% confidence intervals of the novelty contrast. **(B)** Parameter estimates and 90% confidence intervals of the memory contrast. There were no age group differences for any of the scores. This figure mirrors Figure 3 from the main paper.

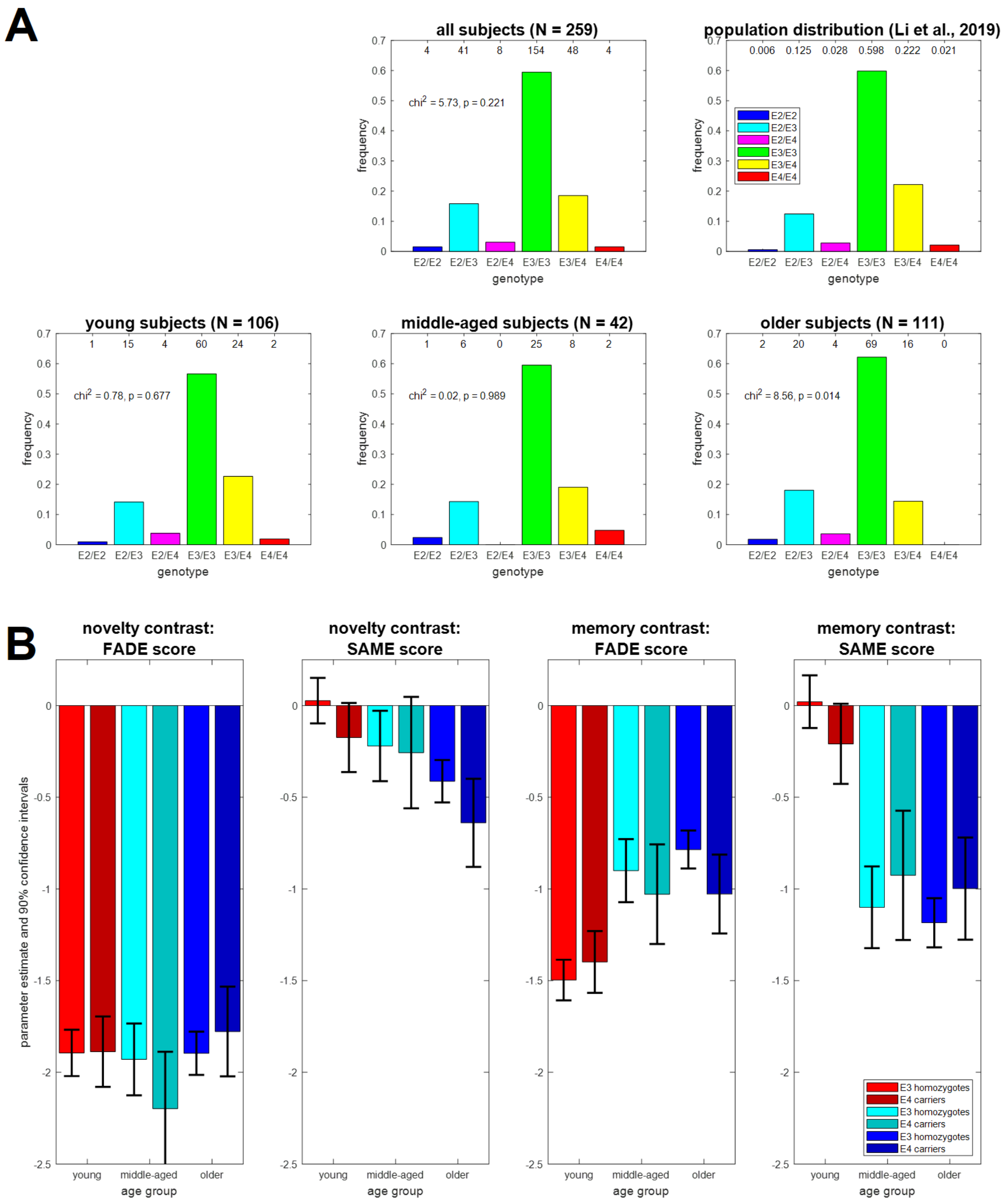


**Figure S3.** *Analysis of FADE scores as a function of ApoE genotype.* **(A)** Distribution of ApoE genotypes in all subjects (top-center) and by age group (bottom row). Frequencies of the population distribution (top-right) were obtained from a behavioral genetics study in a comparable German sample (X. Li et al., 2019). When compared against the population distribution using a chi-squared goodness-of-fit test (df = 2), a significant deviation was observed in older subjects (bottom-right), but not in any of the other groups. **(B)** Results of two-way ANOVAs with age group (young, middle-aged, older) and ApoE genotype (E3 homozygotes: E3/E3; E4 carriers: E3/E4 or E4/E4) as between-subject factors. There was no main effect of or interaction with ApoE genotype (all $p > 0.05$) on any of the scores (FADE-classic, FADE-SAME), computed from any of the contrasts (novelty, memory).

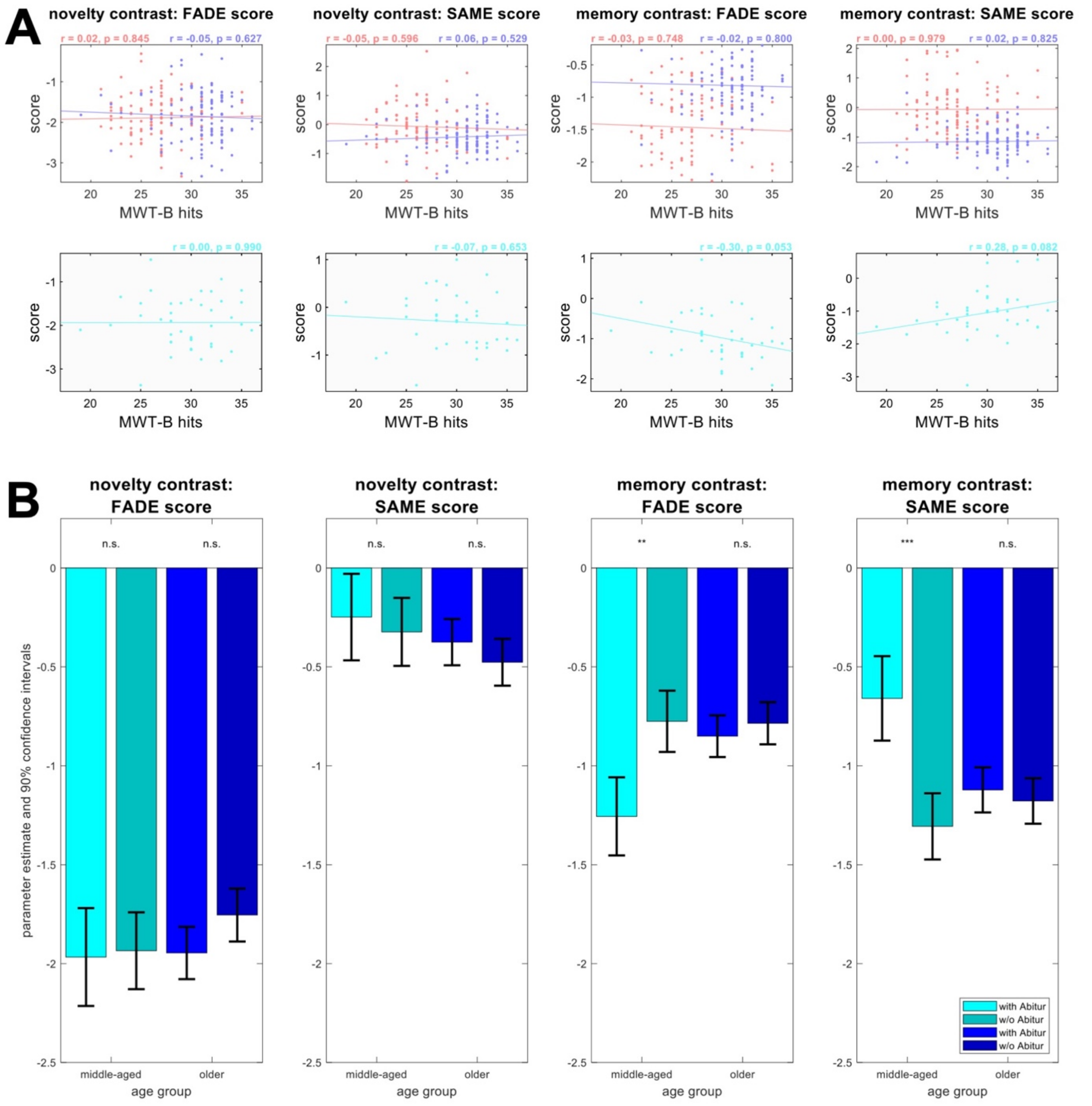


**Figure S4.** *Analysis of FADE scores as a function of educational status.* **(A)** Relationship of FADE-classic and FADE-SAME scores with hits in the MWT-B, a German language vocabulary test (see Section 2.1 and Table 1). There were no significant correlations for young and older adults (top row) or middle-aged subjects (bottom row). **(B)** Results of two-way ANOVAs with age group (middle-aged, older) and educational status (with vs. without Abitur, see Tables 1 and S1) as between-subject factors. Young subjects were excluded from this analysis due to high imbalance with respect to educational status (100 with vs. 6 without Abitur). There were main effects of educational status and interactions with age group on the memory contrast (all $p < 0.05$). Results of within-group two-sample t-tests are reported above the horizontal axis (n.s. = not significant, * $p < 0.05$, ** $p < 0.025$, *** $p < 0.00625$).

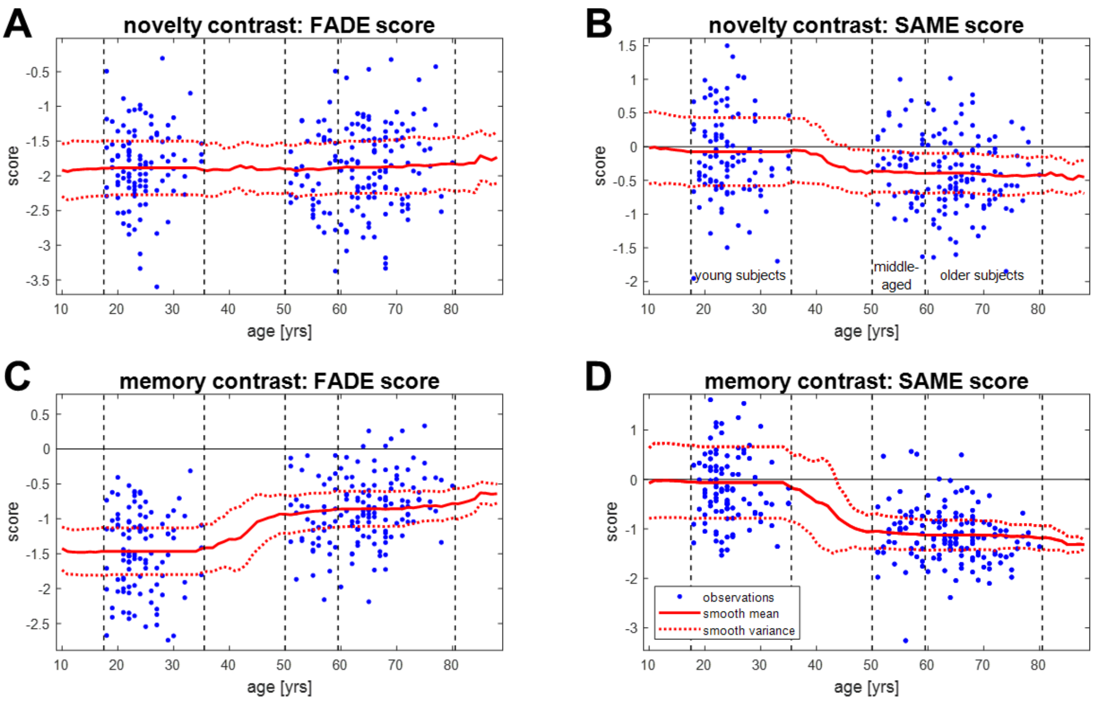


**Figure S5.** *Analysis of FADE scores as a continuous function of age.* FADE scores were plotted against age (blue) and smoothed using a sliding window of 32 years (red). The solid horizontal black line in each panel represents zero, and the dashed vertical black lines in each panel represent limits of age groups (young: 18-35 yrs; middle-aged: 51-59 yrs, older: 60-80 yrs). **(A)** Classic FADE score computed based on the novelty contrast. **(B)** FADE-SAME score computed based on the novelty contrast. **(C)** Classic FADE score computed based on the memory contrast. **(D)** FADE-SAME score computed based on the memory contrast. This figure partly illustrates correlations reported in Figure 4 of the main paper.

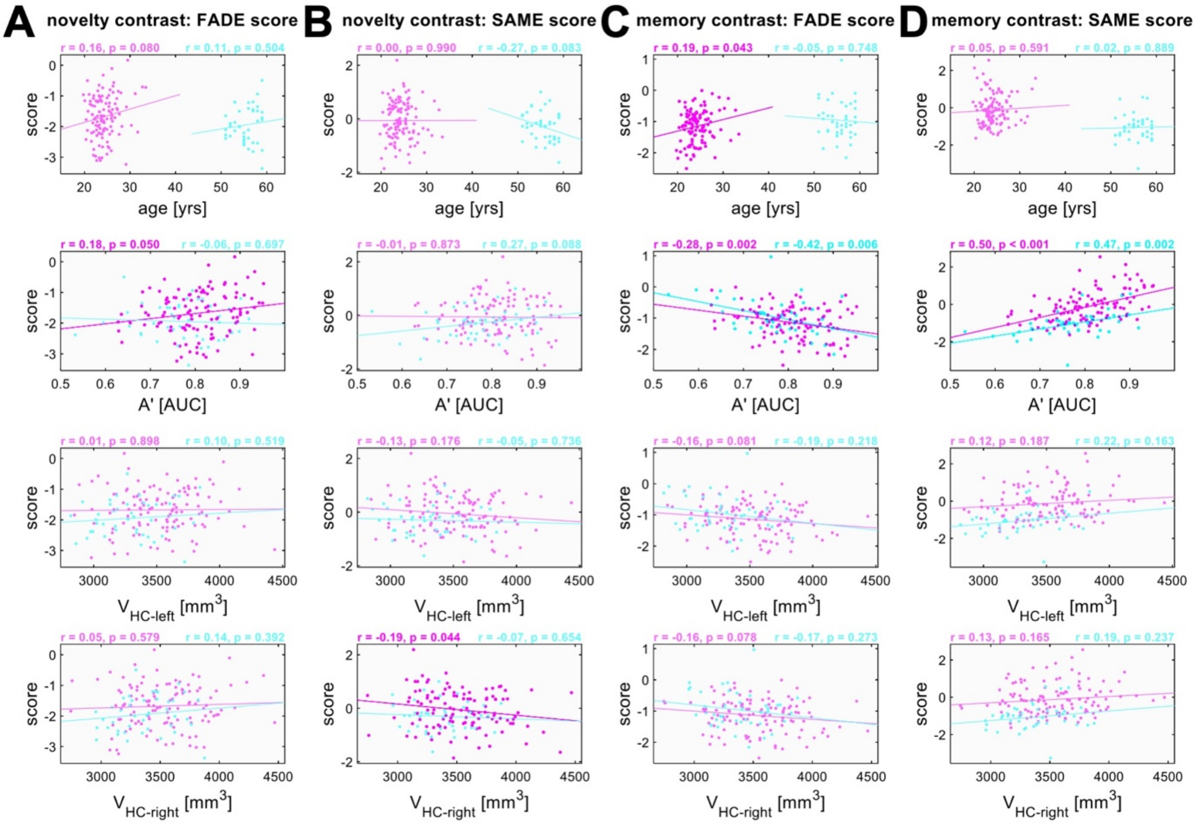


**Figure S6.** *Correlations with independent variables, separated by age group.* Results from correlation analyses of FADE-classic and FADE-SAME scores with age, memory performance (A') and hippocampal volumes ($V_{HC}$). Correlations are reported separately for **(A)** the classic FADE score computed from the novelty contrast, **(B)** the FADE-SAME score computed from the novelty contrast, **(C)** the classic FADE score computed from the memory contrast and **(D)** the FADE-SAME score computed from the memory contrast. Replication subjects are depicted in magenta, and middle-aged subjects are depicted in cyan. Significant correlation coefficients are highlighted. This figure mirrors Figure 4 from the main paper.

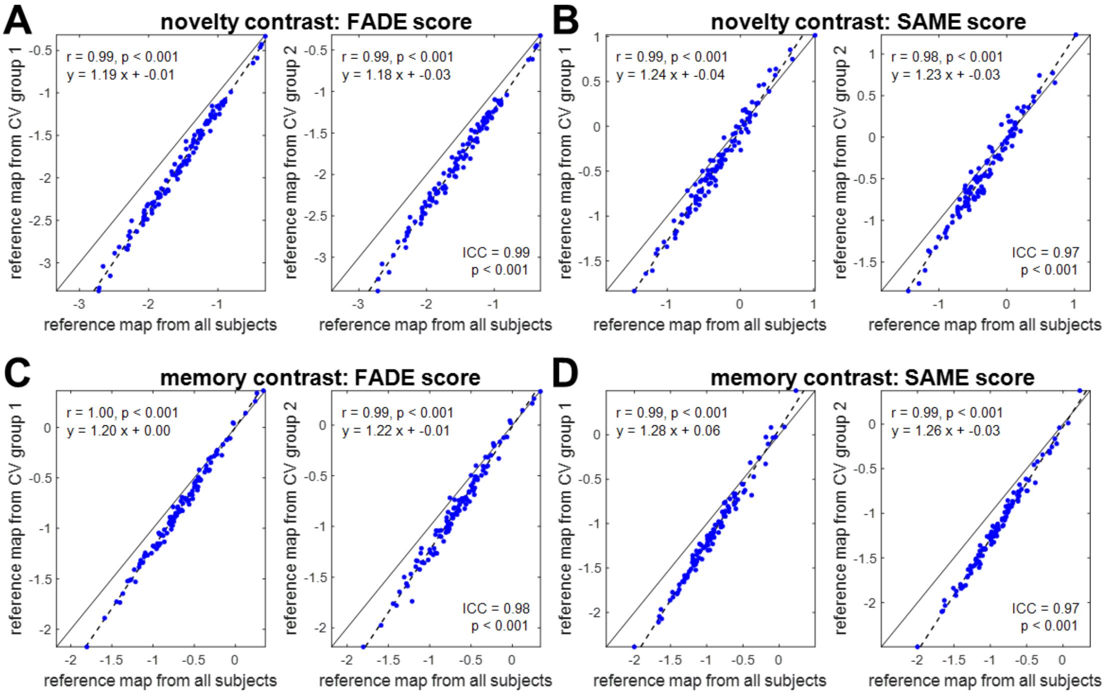


**Figure S7.** *Extraction of FADE scores from all vs. half of young subjects.* Comparison of scores computed for older subjects (older AiA), using reference maps obtained from either all young subjects (young AiA), the first half of those subjects (CV group 1) or the second half of those subjects (CV group 2). In all panels, the solid black line represents the identity function and the dashed black line depicts the regression line. **(A)** Classic FADE score computed based on the novelty contrast. **(B)** FADE-SAME score computed based on the novelty contrast. **(C)** Classic FADE score computed based on the memory contrast. **(D)** FADE-SAME score computed based on the memory contrast. In all comparisons, the correlation coefficient was very high, the intercept of the regression line was approximately zero, and the slope of the regression line was larger than one. This figure extends analyses reported in Figure 6 of the main paper.

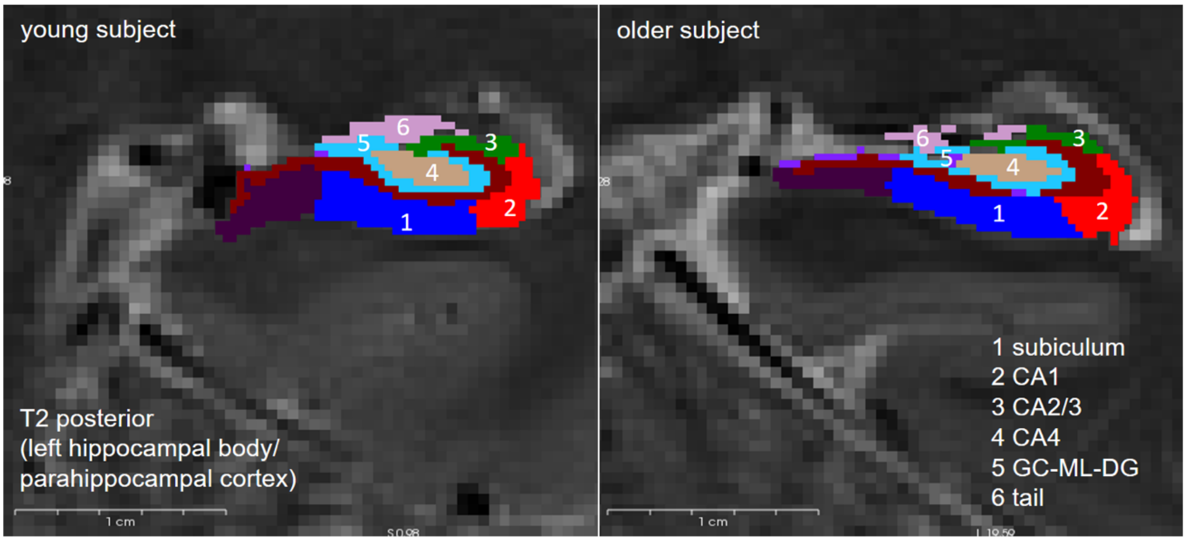


**Figure S8.** *Examples of automatic hippocampal segmentation.* Segmentation results obtained with FreeSurfer superimposed on high-resolution T2-weighted MR images are shown for a representative young subject (left) and a representative older subject (right).
